# Lipopolysaccharide Induces Trained Innate Immune Tolerance in the Heart Through Interferon Signaling in a Model of Stress-Induced Cardiomyopathy

**DOI:** 10.1101/2024.09.24.614798

**Authors:** Kenji Rowel Q. Lim, Junedh Amrute, Attila Kovacs, Abhinav Diwan, David L. Williams, Douglas L. Mann

## Abstract

**Background:** Although the ability of the heart to adapt to environmental stress has been studied extensively, the molecular and cellular mechanisms responsible for cardioprotection are not yet fully understood.

**Methods:** We administered Toll-like receptor (TLR) agonists or a diluent to wild-type mice and assessed their potential to induce cardiac protection against injury from a high intraperitoneal dose of isoproterenol (ISO) administered 7 days later. Cardioprotective effects were analyzed through serum cardiac troponin I levels, immune cell profiling via flow cytometry, echocardiography, and multiomic single-nuclei RNA and ATAC sequencing.

**Results:** Pretreatment with the TLR4 agonist lipopolysaccharide (LPS), but not TLR1/2 or TLR3 agonists, conferred cardioprotection against ISO, as demonstrated by reduced cardiac troponin I leakage, decreased inflammation, preservation of cardiac structure and function, and improved survival. Remarkably, LPS-induced tolerance was reversed by β-glucan treatment. Multiomic analysis showed that LPS-tolerized hearts had greater chromatin accessibility and upregulated gene expression compared to hearts treated with LPS and β-glucan (reverse-tolerized). The LPS tolerance was associated with upregulation of interferon response pathways across various cell types, including cardiac myocytes and stromal cells. Blocking both type 1 and type 2 interferon signaling eliminated LPS-induced tolerance against ISO, while pretreatment with recombinant type 1 and 2 interferons conferred cardiac protection. Multiomic sequencing further revealed enhanced cytoprotective signaling in interferon-treated hearts. Analysis of cell-cell communication networks indicated increased autocrine signaling by cardiac myocytes, as well as greater paracrine signaling between stromal cells and myeloid cells, in LPS-tolerized versus reverse-tolerized hearts.

**Conclusions:** LPS pretreatment confers cardiac protection against ISO-induced injury through TLR4 mediated type 1 and 2 interferon signaling, consistent with trained innate immune tolerance. The observation that LPS-induced protection in cardiac myocytes involves both cell-autonomous and non-cell-autonomous mechanisms underscores the complexity of innate immune tolerance in the heart, warranting further investigation into this cardioprotective phenotype.

**Clinical Perspective:** *What is new?:*

- The Toll-like receptor 4 (TLR4) agonist lipopolysaccharide (LPS) confers cardiac protection against isoproterenol-mediated injury in a manner consistent with trained innate immune tolerance, which is reversed by β-glucan treatment.
- Activation of type 1 and 2 interferon signaling, which is downstream of Toll-like receptor 4, is necessary and sufficient for LPS-induced cardiac protection.
- LPS-tolerized hearts show heightened autocrine signaling by cardiac myocytes and, to a greater degree, increased cell-cell communication between cardiac myocytes and stromal and myeloid cells compared to reverse-tolerized hearts.

*What are the clinical implications?:*

- TLR4 and interferon signaling play key roles in the establishment of cardiac protection and LPS-induced trained innate immune tolerance.
- The protective effects of LPS are mediated by cell-autonomous and non-cell-autonomous mechanisms, suggesting that a deeper understanding of the molecular and cellular signatures of innate immune tolerance is required for the development of targeted approaches to modulate trained innate immunity, and consequently cytoprotection, in the heart.

## Introduction

The ability of the myocardium to adapt to environmental stressors, such as a hemodynamic overload, myocardial ischemia/infarction, or infection, determines whether the heart will decompensate and fail or whether it will maintain preserved function. Three of the important mechanisms that enable the heart to tolerate environmental stress include cardiac hypertrophy, upregulation of cardiac myocyte cytoprotective responses, and cardiac repair mechanisms.^1^ Although these processes are widely recognized as essential for preserving myocardial homeostasis, the specific molecules that mediate and coordinate the response of the heart to environmental stress, both within individual myocytes and across the entire heart, are not fully understood.^1^

We recently demonstrated that isoproterenol (ISO)-induced tissue injury and inflammation was sufficient to protect the heart from the myopathic effects of a subsequent second exposure to ISO, consistent with ISO-induced pre-conditioning.^2^ This study further demonstrated that pharmacologic depletion of macrophages and dendritic cells attenuated the ISO-induced cytoprotective response, raising the intriguing possibility that immune mediated tissue repair mechanisms might precondition the heart to withstand subsequent stress. Relevant to this discussion, Toll-like receptor (TLR) ligands, particularly TLR4, have been shown to induce a transcriptional shift in macrophages, transitioning them from a pro-inflammatory state to an anti-inflammatory and pro-reparative state. This phenotypic shift is essential for maintaining tissue homeostasis by limiting the collateral damage caused by excessive inflammation.^3^ TLR4-mediated repression of transcription was linked primarily to suppression of NF-κB target genes. In contrast, genes containing interferon regulatory factor (IRF) motifs were more likely to be super-induced in tolerant cells.^3^ Recognizing that ISO-induced necrosis of cardiac myocytes results in the release of cytosolic and nuclear proteins containing damage-associated molecular patterns (DAMPs), which in turn activate signaling through endosomal and cell membrane TLRs,^4^ we hypothesized that TLR-mediated signaling might protect the heart from the deleterious effects of ISO. Here we demonstrate that pretreatment with lipopolysaccharide (LPS), a classic TLR4 ligand, induces cross-tolerance in the heart against the myopathic effects of ISO through interferon type 1/type 2 signaling pathways, consistent with the development of trained innate immune tolerance.

## Methods

### Animals

Female 9-10-week-old wild-type C57BL/6J (#000664) were purchased from The Jackson Laboratory (Bar Harbor, ME) and housed at a pathogen-free facility maintained by the Division of Comparative Medicine at the Washington University School of Medicine. Mice had ad libitum access to standard chow diet and water. All animal experiments were performed following the NIH Guide for the Care and Use of Laboratory Animals, as approved by the Institutional Animal Care and Use Committee, Washington University School of Medicine.

### Isoproterenol-induced cardiac injury

Mice were intraperitoneally (i.p.) injected with 300 mg/kg isoproterenol (ISO; Sigma, St. Louis, MO) dissolved in endotoxin-free phosphate-buffered saline (PBS) to model acute, stress-induced cardiac injury, as previously described.^5^ ISO solutions were prepared immediately before injections and kept on ice until use.

### Reagents

To screen for potential TLR ligands that protected against ISO-induced cardiac injury, mice were pretreated with diluent (PBS), 2 mg/kg Pam3CSK4 (#tlrl-pms, InvivoGen, San Diego, CA), 2.5 mg/kg LPS (Sigma), 2.5 mg/kg ultrapure LPS (#tlrl-3pelps, InvivoGen), or 12.5 mg/kg high molecular weight Poly(I:C) (#tlrl-pic, InvivoGen). Mice were also treated with 1 μg recombinant murine interferon-β1 (#581304, Biolegend, San Diego, CA), 10 μg recombinant murine interferon-γ (#575308, Biolegend), a combination of both recombinant murine interferon-β1 and -γ, or diluent (PBS), 500 μg of monoclonal InVivoMAb antibodies against mouse IFNAR-1 (#BE0241, Bio X Cell, Lebanon, NH) or IFNγR (#BE0029, Bio X Cell), their corresponding isotype controls (#BE0083 or #BE0089, respectively, Bio X Cell), and 1 mg of *C. albicans* β-glucan/mouse (kindly provided by Dr. David Williams, East Tennessee State University) or the corresponding vehicle control (5% dextrose in water).

### 2-D Echocardiographic Studies

#### Image acquisition

Ultrasound examination of the cardiovascular system was performed using the Vevo 3100 system (VisualSonics, Toronto, Canada) equipped with an MX400 transducer, as described previously.^5, 6^

#### Imaging protocol

Mice were imaged by echocardiography at baseline, 1 hour, and 6 hours after ISO-injection to evaluate left ventricle (LV) regional and global structure and function, as described.^5^ Avertin (0.005 mL/g) was used for sedation for all imaging studies. Vevo Lab 5.8.1 enabled for the VevoStrain speckle-tracking software (VisualSonics) was used for image analysis of global LV structure and function.

#### Assessment of LV regional wall motion

LV regional wall motion was assessed using a bull’s eye plot that was configured to display 84 segments (7 myocardial cross-sections x 12 radial sections within each cross-section), where the inner ring represents the apex of the LV, the middle ring represents the segments of the mid-LV and the outer ring represents the basal segments of the LV, as described.^5^ LV wall motion within each segment was assessed visually by an experienced operator blinded to the experimental protocol at the time of analysis. Wall motion within each segment was scored as normal, hypokinetic or akinetic. The Segmental Wall Motion Score Index (SWMSI) was then calculated by summing the wall motion scores for all LV segments and dividing by the total number of segments in the LV wall. A value greater than 1 indicates abnormal segmental wall motion, as previously described.^5^

### Serum Troponin I Release

Blood was collected from mice by puncturing the submandibular vein with an 18G needle and letting the blood drip into BD Microtainer^TM^ tubes (BD Biosciences, San Jose, CA). Samples were spun at 1,000 rcf for 5 min at room temperature, after which the sera were collected, diluted 1:4 with PBS, and analyzed with the ARCHITECT i2000 analyzer (Abbott Laboratories, Abbott Park, IL).

### Flow cytometry

The enumeration of immune cells in murine tissues through flow cytometry was performed as previously described.^5, 7^ Blood was collected through submandibular vein puncture into EDTA-coated collection tubes, after which the mice were euthanized by CO_2_ and cervical dislocation. Hearts were perfused with ice-cold Hanks’ balanced salt solution (HBSS, no calcium/magnesium, Corning, Glendale, AZ) to drain off excess blood, and then isolated with the atria and connected vessels removed. Hearts were weighed, minced, and digested with 450 U/mL collagenase (Sigma), 60 U/mL DNase I (Sigma), and 60 U/mL hyaluronidase (Sigma) in HBSS for 45 min at 37°C, with shaking. HBSS with 2% fetal bovine serum (Thermo Fisher Scientific, Waltham, MA) and 0.2% bovine serum albumin (HBB solution) was then added to the mixtures to halt enzymatic digestion. Spleens were isolated and smashed with the flat end of a syringe in ice-cold HBB. Tibias were isolated, and their lengths were measured to calculate for the heart weight-to-tibia length ratio. Heart and spleen cell suspensions were strained through a 40 μm filter, and spun at 350 rcf, 4°C for 5-7 min. The cell pellets and 20 μL of blood were then dissolved/incubated in ACK lysing buffer (Thermo Fisher Scientific) for 15 min on ice. The suspensions were subsequently diluted with flow cytometry buffer (2% fetal bovine serum and 2 mM EDTA in PBS), and spun at 350 rcf, 4°C for 5 min. Finally, the cell pellets were dissolved/incubated in flow cytometry buffer containing fluorescent-tagged antibodies diluted 1:400 from stock for 30 min at 4°C. The antibodies used were: CD45-PerCp/Cy5.5, Ly6G-FITC, CD11b-BV510, CD64-PE, Ly6C-APC/Cy7, CD19-APC, CD4-PE/Cy7 (Biolegend), and CD8a-BV421 (BD Biosciences). Excess antibody solution was removed and replaced with flow cytometry buffer, and cells were analyzed using the BD LSRFortessa or X-20 (BD Biosciences) at the Flow Cytometry & Fluorescence Activated Cell Sorting Core, Department of Pathology and Immunology, Washington University School of Medicine. Compensation controls were created using UltraComp eBeads (Invitrogen, Carlsbad, CA). Counts are expressed as the number of cells per mg tissue for the heart, the number of cells per 20 μL volume for the blood, or as the percentage of CD45^+^ cells for the spleen (at least 400,000 total cells collected). The gating strategy used is shown in **Supplemental Figure S1**.

### Gravimetric and Histological Analysis

Mice were euthanized at baseline, and 1, 3 and 7 days after ISO-injection, and the hearts were removed and weighed to determine the heart weight/tibia length (HW/TL) ratio. Mouse hearts were perfused with ice-cold PBS, dissected out, weighed, and transversely cut in half between the base and the apex. The apical half was incubated overnight in 10% formalin at 4°C, followed by three 10-min washes in PBS, and consecutive 30-min incubations in 20%, 50%, and 70% ethanol. Fixed hearts were maintained in 70% ethanol and submitted for paraffin-embedding and stained with hematoxylin and eosin and Masson’s trichrome in the Advanced Imaging and Tissue Analysis Core, Digestive Disease Research Core Center, Washington University School of Medicine. Samples were visualized using the Zeiss Axioskop brightfield microscope (Zeiss, Oberkochen, Germany), taking images of 6 random fields of view for each sample. The degree of inflammation in each hematoxylin/eosin-stained image was then visually scored according to the following criteria: 0, no infiltrate; 1, <25% infiltration; 2, 25-50% infiltration; 3, 50-75% infiltration; and 4, 75-100% infiltration, as previously described.^5^

### snRNA-seq and snATAC-seq

Mice were treated with LPS (2.5 mg/kg), β-glucan (1 mg/mouse), 1 μg recombinant murine interferon-β1/10 μg recombinant murine interferon-γ or diluent. Mice were euthanized 7 days later (n=3 per group), in which hearts were perfused with ice-cold PBS and dissected out. After removing the atria and connected vessels, hearts were minced and snap-frozen in liquid nitrogen. Frozen hearts were used as starting material for the Chromium Nuclei Isolation kit (10x Genomics, Pleasanton, CA), with isolated nuclei from all hearts belonging to the same group of treated mice pooled together prior to submission to the Genome Access Technology Center at the McDonnell Genome Institute, Washington University in St. Louis for single nuclei multiome ATAC + Gene Expression sequencing.

Raw fastq files were aligned using CellRanger ARC to generate fragment files and aligned count matrices. Data was subsequently analyzed in the Seurat and Signac framework. First peaks from individual sample bed files between 20-10,000 bp were kept. Then a merged Seurat object was created using a common peak set. We then applied QC filters for the nuclei to only include those satisfying nCount_ATAC < 25000, nCount_RNA < 25000, nCount_ATAC > 500, nCount_RNA > 500, nucleosome_signal < 2, TSS.enrichment > 1, and percent.mt < 10. Peaks were called using MACS2. LSI was used for ATAC and PCA for RNA for dimensional reduction. FindMultiModalNeighbors in Seurat was then used to build a joint neighbor graph using both assays which was then used for UMAP embedding construction. We used several clustering resolutions and DE analysis to annotate nuclei clusters based on canonical RNA marker genes. DE RNA analysis was performed in each cell type using FindAllMarkers and genes with p-adj < 0.05 and log2FC > 0.25 were deemed significant. Differential peaks were calculated using FindMarkers with the LR test and nCount_peaks as the latent variable. Peaks were deemed significant if p-val < 0.005 and avg_log2FC > 0.25 (as per Signac). Peaks were linked to genes using the LinkPeaks function. Pathway enrichment analysis was performed using Enrichr, with the MSigDB Hallmark 2020 database.^8–10^ For heatmaps using differentially expressed genes, gene lists were obtained from the Gene Transcription Regulation Database (GTRD) or the Gene Ontology database (GO) and used to filter our DE RNA dataset.

To dissect cell interaction networks, we used CellChat.^11^ Briefly, we split the data into tolerized (LPS+vehicle) and reverse-tolerized (LPS+β-glucan) nuclei and used the integrated cell annotations to run CellChat using the CellChatDB database ignoring non-protein signaling. To compute network centrality scores, we used netAnalysis_computeCentrality. Finally, to visualize differential signaling in the tolerized and reverse-tolerized states, we merged the CellChat output from the two analyses and used the netVisual_diffInteraction and netVisual_heatmap to visualize high level changes.

### qPCR

Mice were i.p. injected with 2.5 mg/kg LPS, and hearts were collected at baseline, and at days 1 and 7 after treatment. Hearts were perfused with ice-cold PBS and dissected out. The inferior mid-apical portion of the LV for each heart was then isolated and snap-frozen; this part of the heart was selected as the starting material for RNA sequencing as it had the most abnormal wall motion 1 hr post-ISO, as found in our previous study.^5^ Total RNA was extracted using the RNeasy Fibrous Tissue Mini Kit (Qiagen, Germantown, MD) following manufacturer’s instructions, and 1000 ng was used as template for cDNA synthesis with the High-Capacity RNA-to-cDNA Kit (Applied Biosystems, Waltham, MA), also following manufacturer’s instructions. For qPCR, cDNA was diluted at a 1:4 ratio with DNase/RNase-free distilled water, and then mixed with TaqMan Fast Advanced Master Mix (Applied Biosystems) and predesigned TaqMan Gene Expression Assays (Applied Biosystems) for *Ifnb1* (Mm00439546_s1), *Ifna2* (Mm00833961_s1), *Ifit1* (Mm00515153_m1), *Ifng* (Mm01168134_m1), *Irf1* (Mm01288580_m1), or *Gapdh* (Mm99999915_g1) following manufacturer’s instructions. Samples were then run in triplicate on the QuantStudio 3 (Applied Biosystems) using the default Fast program, and the 2^−ΔΔCt^ method was used to calculate relative expression, normalized to *Gapdh* expression. To confirm the modulation of downstream interferon signaling after anti-IFNAR1/IFNγR antibody or recombinant murine interferon treatment, total RNA extraction and qPCR for *Ifit1*, *Irf1*, and *Gapdh* were performed using mouse heart samples as described above.

### Statistical analysis

All data are presented as mean ± standard error of the mean (SEM). The Shapiro-Wilk test was used to determine whether the data were normally distributed. One-way analysis of variance (ANOVA) with Dunnett’s (multiple comparisons to a control) or Tukey’s (all pairwise comparisons) correction for multiple post-hoc comparisons were performed, where appropriate. For regular or repeated measures two-way ANOVA, a Sidak’s correction was used for post-hoc comparisons. Unpaired two-tailed t-tests were performed where appropriate. The Kaplan-Meier survival curves were analyzed using the log-rank (Mantel-Cox) test. For data that were not normally distributed we performed a Kruskal-Wallis test, using Dunn’s post-hoc testing where appropriate, or a Mann-Whitney test. All statistical analysis was performed using GraphPad Prism 9 (GraphPad Software, Boston, MA). A p-value < 0.05 was considered statistically significant.

## Results

### Screening for ligands that confer cytoprotective responses in the heart following ISO-injury

To screen for TLR ligands that conferred cytoprotective responses in the heart following ISO-induced cardiac injury, we pretreated wild-type mice with diluent (PBS), Pam3CSK4 (TLR1/2), poly(I:C) (TLR3), or LPS (TLR4) 7 days prior to ISO (300 mg/kg, i.p.) injection **(Figure 1A)**. TLR ligands were selected based on the pathways that are activated by DAMPs:^12^ TLR1/2 stimulated NF-κB signaling, TLR3 stimulated interferon signaling, and TLR4 stimulated both pathways.^13^ The concentrations of TLR ligands chosen were based on previous studies in which these doses induced cardioprotection against myocardial ischemia/reperfusion injury, as well as through pilot studies.^14–16^ As shown in **Figure 1B**, ISO injection provoked significant increases in troponin I levels in mice pre-treated with diluent (*P*=0.005), Pam3CSK4 (*P*=0.045), or poly(I:C) (*P*=0.044) when compared to baseline values (day 0). Remarkably, ISO injection did not provoke a significant increase in cardiac troponin I levels in mice pre-treated with LPS (*P*>0.99) **(Figure 1B)**, suggesting that TLR4 signaling conferred cytoprotective effects. As an additional method for screening and identifying cytoprotective TLR ligands, we performed flow cytometry at baseline (day 0) and 1 day after ISO injection using the hearts of mice pretreated with PBS, Pam3CSK4, poly(I:C), or LPS. As shown in **Figure 1C**, there were no overall changes in the number of CD45^+^ cells in the heart among the different treatment groups on day 1 after ISO-injection when compared to baseline values. There were, however, significant increases in the number of Ly6G^+^ neutrophils migrating into the hearts on day 1 after ISO-injury in the PBS-(*P*=0.0004) and poly(I:C)-treated (*P*=0.008) mice (**Figure 1D**). Although there were numeric increases in Ly6G^+^ neutrophils in the hearts of the Pam3CSK4 treated mice, these changes were not significant statistically (*P*=0.116). Remarkably, LPS pre-treatment prevented the ISO-injury induced increase in Ly6G^+^ neutrophil influx on day 1 (*P*>0.99 compared to baseline values). Viewed together, these findings suggest that TLR4 signaling confers short-term protective effects in the heart following ISO-induced cardiac injury, whereas activation of TLR1/2 and TL3 signaling does not confer short-term cytoprotective effects.

**Figure 1.**
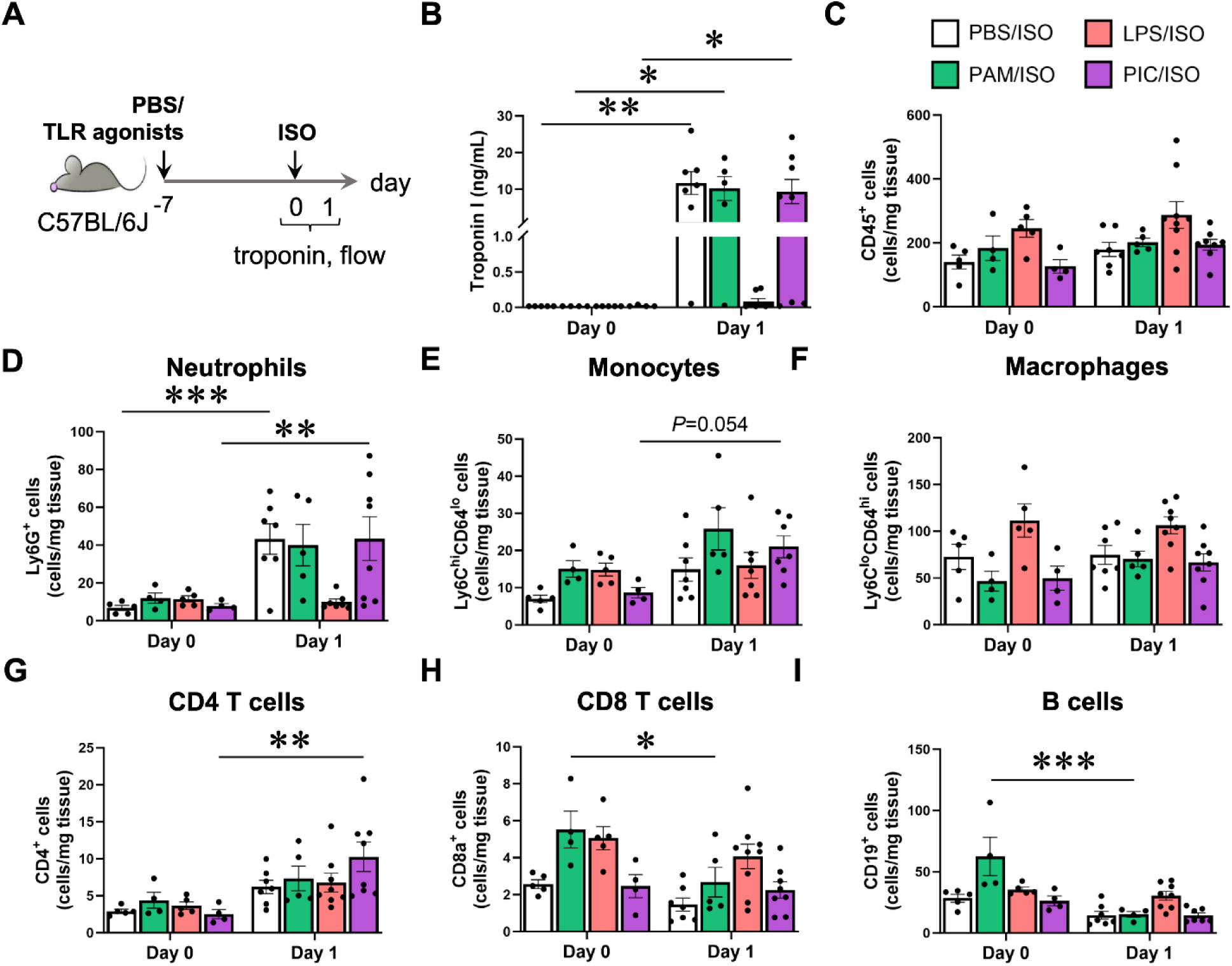
Cardioprotective effects TLR agonists against ISO-injury. **(A)** Mice were injected i.p. with Pam3CSK4 (PAM; 2 mg/kg), lipopolysaccharide (LPS; 2.5 mg/kg), poly(I:C) (PIC; 12.5 mg/kg), or diluent (PBS) on day -7, and challenged with an i.p. injection of ISO (300 mg/kg) on day 0. **(B)** Serum cardiac troponin I analysis on days 0 and 1. Flow cytometry analysis of **(C)** total CD45^+^ cells, **(D)** Ly6G^+^ neutrophils, **(E)** Ly6C^hi^CD64^lo^ monocytes, **(F)** Ly6C^lo^CD64^hi^ macrophages, **(G)** CD4^+^ T cells, **(H)** CD8a^+^ T cells, and **(I)** CD19^+^ B cells in the heart on days 0 and 1, with counts expressed as cells/mg of tissue (n=4-9/group). Data show mean ± SEM and were analyzed by two-way ANOVA with Sidak’s multiple comparisons test. (Key: \**P*<0.05, \*\**P*<0.01, \*\*\**P*<0.001)

Recognizing that standard preparations of LPS may contain impurities that activate TLRs other than TLR4,^17, 18^ we repeated the above experiments using an ultrapure TLR4-specific formulation of LPS that did not contain the TLR2-activating lipoproteins found in standard LPS preparations and compared the results with the LPS preparation employed in the TLR screening studies. As shown in **Supplemental Figure S2**, the LPS used in our screen yielded similarly reduced troponin I levels (*P*=0.832) as those obtained with the ultrapure TLR4-specific formulation of LPS on day 1 after ISO-injury. Accordingly, we used the standard LPS preparation for all of the studies presented herein.

### Characterization of the LPS induced cytoprotection against ISO-injury

The above studies suggest that LPS pretreatment confers short-term cardioprotective benefits following ISO-injury. To further characterize the cytoprotective effects of LPS, we pre-treated wild-type (WT) mice with either diluent (PBS) or LPS on day -7, and then treated the mice with ISO on day 0 (**Figure 2A**). As shown in **Figure 2B**, there was only 1 death observed in the LPS pretreated group, whereas there was a significant (*P*=0.049) 30% increase in ISO-induced lethality in the diluent-treated group. No differences in HW/TL were observed between groups across all timepoints examined (at least *P*>0.106) **(Figure 2C)**. To characterize the myocardial inflammatory response to ISO-injury in diluent and LPS pretreated mouse hearts, we performed immunohistochemistry and FACS at baseline (day 0) and on days 1, 3 and 7. **Figure 2D** shows representative hematoxylin and eosin staining of leukocyte infiltrates at baseline and on days 3 and 7 after ISO-injection; the results of the group data are summarized in **Figure 2F**. The inflammatory score index was significantly greater on day 3 and day 7 (*P*<0.0001 for both) in the diluent treated mice, whereas there was a numerically smaller but statistically significant (*P*=0.046) increase in the inflammatory score index on day 3 in the LPS pretreated mice (**Figure 2F**). Masson’s trichrome staining revealed that there was increased patchy fibrosis on days 3 and 7 in the hearts of the diluent pre-treated mice relative to LPS pretreated mice (**Figure 2E**).

**Figure 2.**
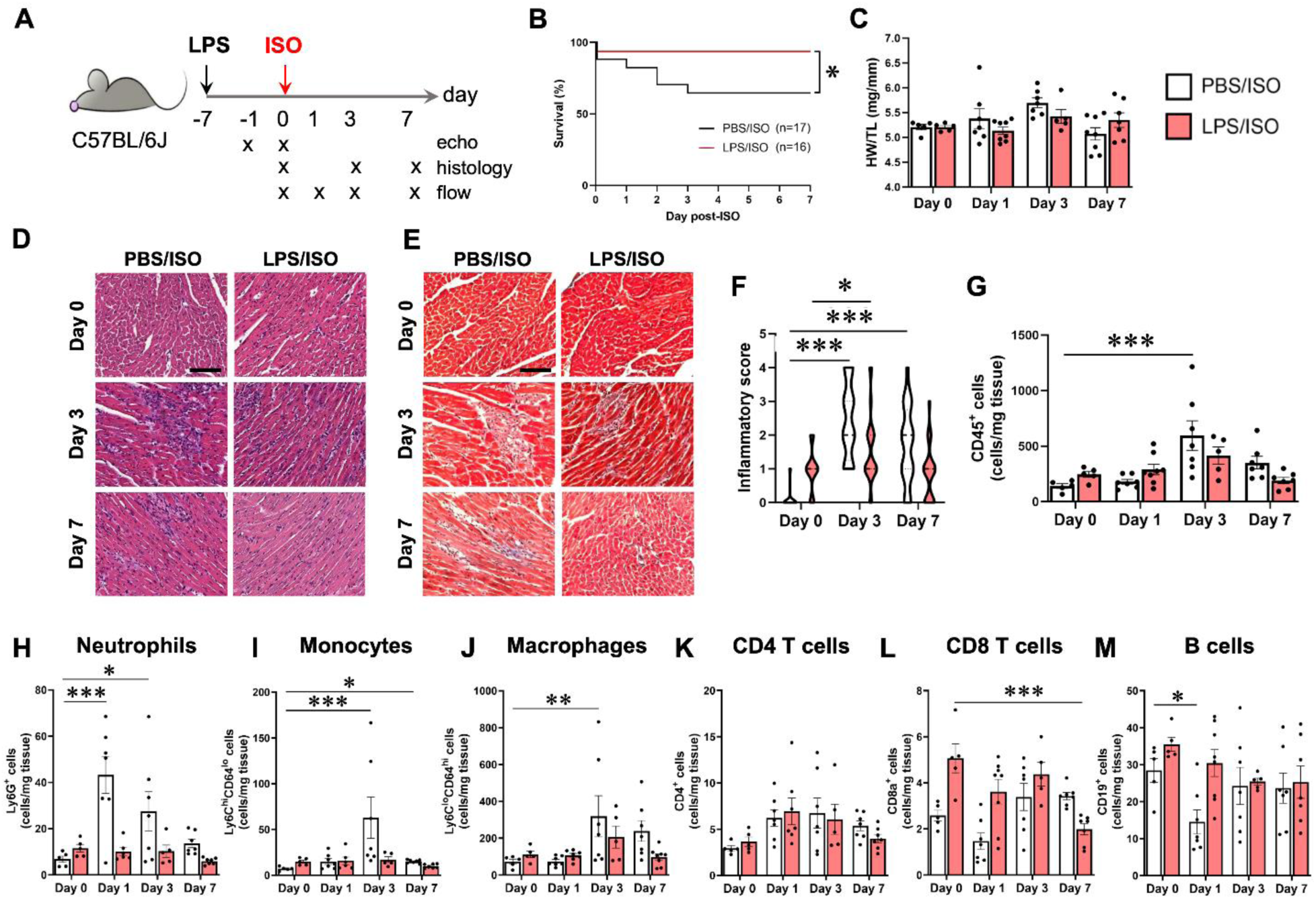
Characterization of LPS pretreated mice following ISO-injury. **(A)** Mice were injected i.p. with lipopolysaccharide (LPS; 2.5 mg/kg) or saline (PBS) on day -7 and challenged with an i.p. injection of ISO (300 mg/kg) on day 0. PBS/ISO and LPS/ISO mice were evaluated as indicated. **(B)** Survival curves until day 7 post-ISO (n=16-17/group). **(C)** Heart weight-to-tibia length (HW/TL) ratios on days 0, 1, 3, and 7 (n=5-8/group). **(D)** Hematoxylin/eosin and **(E)** Masson’s trichrome staining of hearts on days 0, 3, and 7. (Scale bar: 100 μm). **(F)** Quantification of inflammatory scores based on hematoxylin/eosin-stained heart images on days 0, 3, and 7 (n=24 fields/group/time, from 3 hearts/group/time). Flow cytometry analysis of **(G)** total CD45^+^ cells, **(H)** Ly6G^+^ neutrophils, **(I)** Ly6C^hi^CD64^lo^ monocytes, **(J)** Ly6C^lo^CD64^hi^ macrophages, **(K)** CD4^+^ T cells, **(L)** CD8a^+^ T cells, and **(M)** CD19^+^ B cells in the heart on days 0, 1, 3, and 7 (n=5-8/group). Data show mean ± SEM and were analyzed by the log-rank Mantel-Cox test (B), multiple Mann-Whitney test (C), or two-way ANOVA with Dunnett’s multiple comparisons test (F-M). Data for days 0 and 1 were from the same mice used in Figure 1. (Key: \**P*<0.05, \*\**P*<0.01, \*\*\**P*<0.001)

### Changes in cell profiles of cardiac resident immune cells

To gain further insights into the cytoprotective effects of LPS, we evaluated the time-dependent numerical changes in cardiac-resident immune cells, in diluent and LPS pre-treated mice at baseline and on days 1, 3 and 7 after ISO-injection. FACS analysis of the heart showed that relative to baseline values there was a significant increase in CD45^+^ leukocytes in the hearts of PBS-treated mice on day 3 (*P*<0.0001), whereas there was no significant increase in the numbers of CD45^+^ leukocytes in the LPS pretreated hearts at any time point examined (**Figure 2G)**. As shown in **Figure 2H**, the number of Ly6G^+^ neutrophils peaked in the hearts of diluent pretreated mice on day 1 (*P*<0.0001) after ISO-injury and remained significantly elevated above baseline levels on day 3 (*P*=0.019), returning to baseline values on day 7 (*P*=0.73 relative to baseline). The numbers of Ly6C^hi^CD64^lo^ monocytes and Ly6C^lo^CD64^hi^ macrophages in the heart peaked on day 3 (*P*<0.0001 and *P*=0.008, respectively) after ISO-injury; the monocyte counts remained slightly increased on day 7 (*P*=0.030, relative to baseline) while the macrophage counts returned to baseline values on day 7 (*P*=0.093, relative to baseline) (**Figures 2I, 2J**). Remarkably, there was no ISO-injury induced increase in the numbers of Ly6G^+^ neutrophils, Ly6C^hi^CD64^lo^ monocytes, and Ly6C^lo^CD64^hi^ macrophages in the LPS pre-treated mouse hearts at any of the times examined (**Figure 2H-2J)**. While ISO-injury did not induce any significant changes in the CD4^+^ T cell count in either group **(Figure 2K)**, we did observe a significant decrease (*P*<0.0001) in the numbers of CD8^+^ T cells in the hearts of LPS pretreated mice on day 7 after ISO-injury **(Figure 2L)**. Moreover, diluent-treated hearts showed a significant reduction in CD19+ B cells only at day 1 after ISO-injury, whereas LPS pretreated hearts showed no such change **(Figure 2M)**.

### Changes in cell profiles of circulating immune cells

The changes in the number of circulating immune cell subsets in the diluent and LPS pretreated mice at baseline and on days 1, 3 and 7 following ISO-injection are summarized in **Supplemental Figures S3A-F**. Similar to the observations in the heart, when compared to baseline there was a significant ISO-induced (*P*=0.0002) increase in circulating Ly6G^+^ neutrophils on day 1 after ISO in the diluent but not the LPS pretreated mice (*P*=0.535). In contrast to the observations in the heart, there were significant decreases in the number of circulating CD45^+^ leukocytes (*P*=0.015 on day 3), CD4^+^ (*P*=0.006 on day 1, *P*=0.0001 on day 3, *P*=0.001 on day 7) and CD8^+^ T (*P*=0.002 on day 3) cells, as well as CD19^+^ B lymphocytes (*P*<0.0001 on days 1 and 3, *P*=0.0001 on day 7) in the diluent-treated mice, whereas the only decrease in leukocyte population subsets observed in the LPS pre-treated mice was a significant decrease in Ly6C^hi^CD64^lo^ monocytes observed on day 1 (*P*=0.001) and day 7 (*P*=0.008), and in CD19^+^ B cells on day 3 (*P*=0.010).

### Changes in cell profiles of splenic immune cells

The changes in the immune cell profile of splenic cells, expressed as the % of the total number of CD45^+^ cells at baseline and on days 1, 3 and 7 following ISO-injection are summarized in **Supplemental Figures S3G-L**. Relative to baseline values, the changes in the % of immune cell subsets in the spleen were qualitatively similar to those observed in the circulation and heart, with respect to the significant increase (*P*<0.0001) in the % of Ly6G^+^ neutrophils on day 1 and significant decreases in CD19^+^ B lymphocytes on days 1 (*P*=0.0004) and 7 (*P*<0.0001) in the diluent-treated mice. Of note there was a significant increase in the % of splenic CD8^+^ T cells in the diluent-treated mice on day 7 (*P*<0.0001). In contrast to the ISO-induced changes observed in the heart and the circulation following LPS pretreatment, there was a small but significant (*P*=0.034) decrease in the % of CD4^+^ T cells in the spleen on day 7 and a trend towards a significant decrease in the % of splenic Ly6C^hi^CD64^lo^ monocytes on day 1 and day 7 (*P*=0.056 for both).

### Effects of LPS on LV structure and function

To ascertain whether the ISO-induced tissue injury resulted in changes in LV structure and function in the diluent and LPS pretreated mouse hearts, we performed 2-D echocardiography at baseline and at 2 sequential time points after ISO-injection. **Figure 3A** shows bull’s eye plots for diluent and LPS pretreated hearts, **Figure 3B** summarizes the changes in segmental wall motion in different myocardial segments, whereas the group data are summarized in **Figures 3C-3E**. As shown in **Figure 3A-3B**, relative to baseline values, ISO-injection provoked segmental wall motion abnormalities in the mid to apical inferior wall of the LV that were detectable as early as 1 hour after ISO-injection (*P*<0.0001 compared to baseline) in the diluent treated hearts and persisted as long as 6 hours (*P*<0.0001 compared to baseline), consistent with our prior observations.^5^ In contrast, ISO-injection did not provoke significant segmental wall motion abnormalities in the LPS pretreated mice. The overall SWMSI score was significantly greater (*P*=0.036) at 1 hr in the diluent-treated mice when compared to LPS pretreated mice **(Figure 3C)**. We also observed overall significant differences in the LV ejection fraction (LVEF) between diluent-treated and LPS pretreated mice (*P*=0.032, by two-way ANOVA) following ISO-injection, with an ∼8% decrease in LVEF at 1 hour in the diluent-treated mice relative to baseline as opposed to a ∼2% decrease in LPS pretreated mice **(Figure 3D)**. Lastly, there were no significant differences in the LV end-diastolic volume (LVEDV) between diluent-treated and LPS pre-treated mice following ISO-injection at any time point examined **(Figure 3E)**.

**Figure 3.**
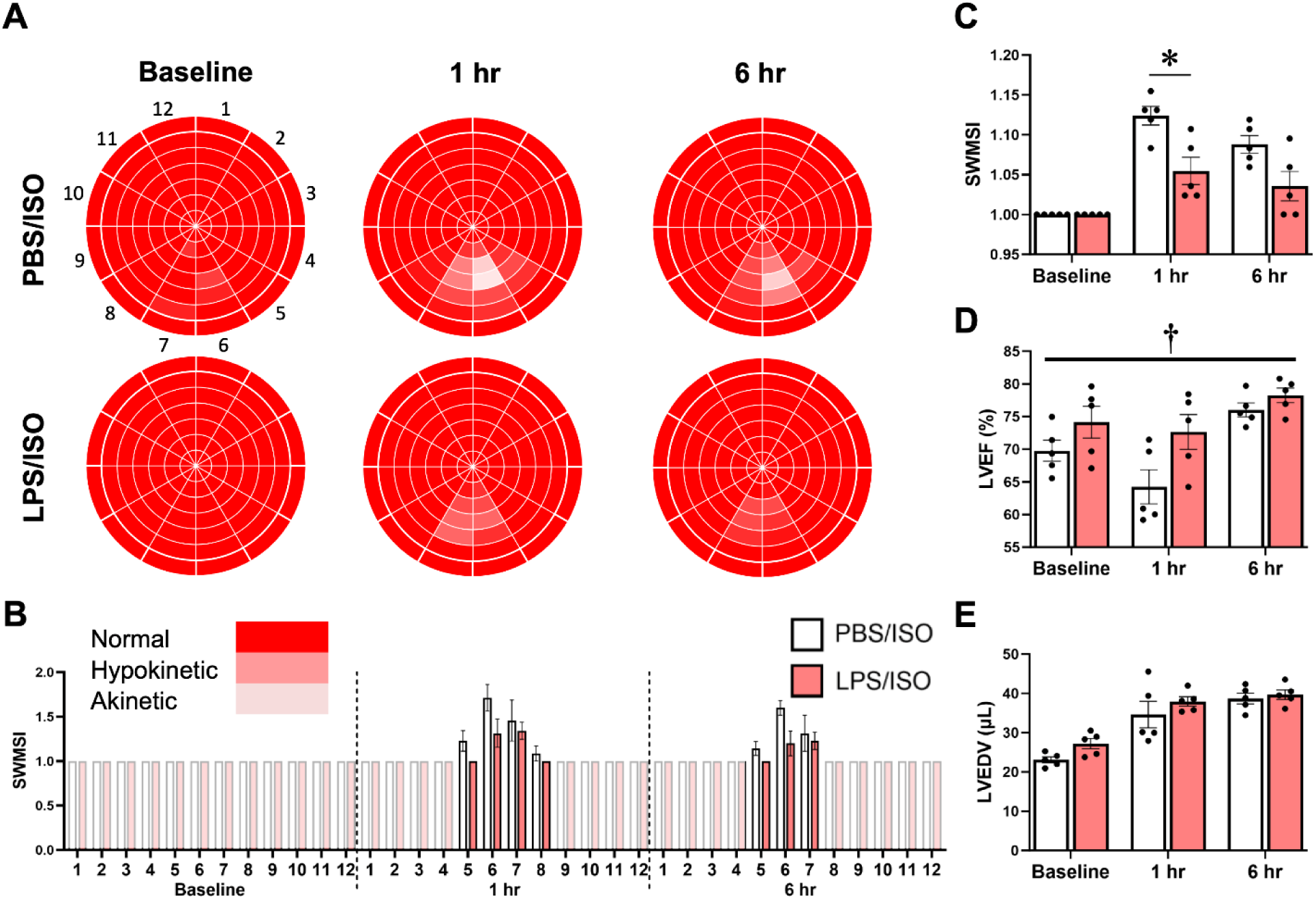
Evaluation of left ventricular structure, function and regional wall motion. Mice were injected i.p. with lipopolysaccharide (LPS; 2.5 mg/kg) or saline (PBS) on day -7 and challenged with an i.p. injection of ISO (300 mg/kg) on day 0. PBS/ISO and LPS/ISO mice were evaluated by 2-D echocardiography at baseline, 1 hr, and 6 hr after ISO injection **(A)** Bulls eye plots showing regional wall motion across the left ventricle with 7 myocardial cross-sections (each containing 12 radial segments) from base to apex. **(B)** Quantification of the segmental wall motion score index (SWSMI) per segment in (A). **(C)** The global SWMSI was determined as the average of 84 regional LV segments, where 1 = normal wall thickening, 2 = hypokinesis, 3 = akinesis **(D)** Left ventricular ejection fraction (LVEF), and **(E)** left ventricular end-diastolic volume (LVEDV) (n=5/group). Data show mean ± SEM and were analyzed by repeated measures two-way ANOVA with Sidak’s multiple comparisons test (C-E). (Key: \**P*<0.05, ^†^ P<0.05 by two-way ANOVA).

### LPS-induced tolerance against ISO-injury is abolished by β-glucan

The finding that pretreatment with LPS seven days prior to ISO injection protected against ISO-induced myocyte injury, LV dysfunction, myocardial inflammation, and increased mortality suggested that LPS exposure resulted in reduced sensitivity (i.e., tolerance) to ISO-mediated cardiac injury. Based on prior studies that showed that LPS tolerance can be mediated through epigenetic changes and that LPS-induced epigenetic programming in immune cells can be reversed by β-glucan,^19^ we asked whether treatment with β-glucan was sufficient to reverse the cytoprotective effects conferred by LPS. For these experiments, mice were treated with LPS on day -7 and were then given β-glucan or vehicle on day -6 followed by ISO-injection on day 0 **(Figure 4A)**. Remarkably, there was a significant (*P*=0.019) ISO-induced increase in serum cardiac troponin I levels (day 1 post-ISO) in the β-glucan-treated mice, whereas troponin I levels were not detectable following ISO-injection in vehicle-treated mice **(Figure 4B)**. There were, however, no significant differences between β-glucan-treated and vehicle-treated mice with respect to the HW/TL ratio **(Figure 4B)** or the number of myocardial CD45^+^ cells in the heart **(Figure 4C**). Flow cytometry analysis of myocardial immune cell subsets showed that there was a significant increase in the number of Ly6G^+^ neutrophils (*P*=0.033) and decrease in the number of Ly6C^lo^CD64^hi^ macrophages (*P*=0.036) and CD19^+^ B cells (*P*=0.022) in the heart (day 1 post-ISO) in the mice treated with β-glucan when compared to vehicle-treated controls **(Figure 4D)**, consistent with a loss of LPS-induced tolerance against ISO-injury. β-glucan treatment also partially reversed the LPS-induced cardioprotective effects on LV cardiac structure and function. As shown in **Figures 4E-4G**, there was a significant (*P*=0.015) increase in ISO-induced segmental wall motion abnormalities in the inferior mid-apical region of the LV in β-glucan-treated mice at 1 hour when compared to vehicle-treated controls. No significant differences between groups were observed for the LVEF (*P*=0.573, two-way ANOVA) **(Figure 4H)** and the LVEDV (*P*=0.717, two-way ANOVA) at 1- and 6-hours post-ISO **(Figure 4I)**. Viewed together, these results demonstrate that treatment with β-glucan reversed LPS-induced tolerance against ISO injury.

**Figure 4.**
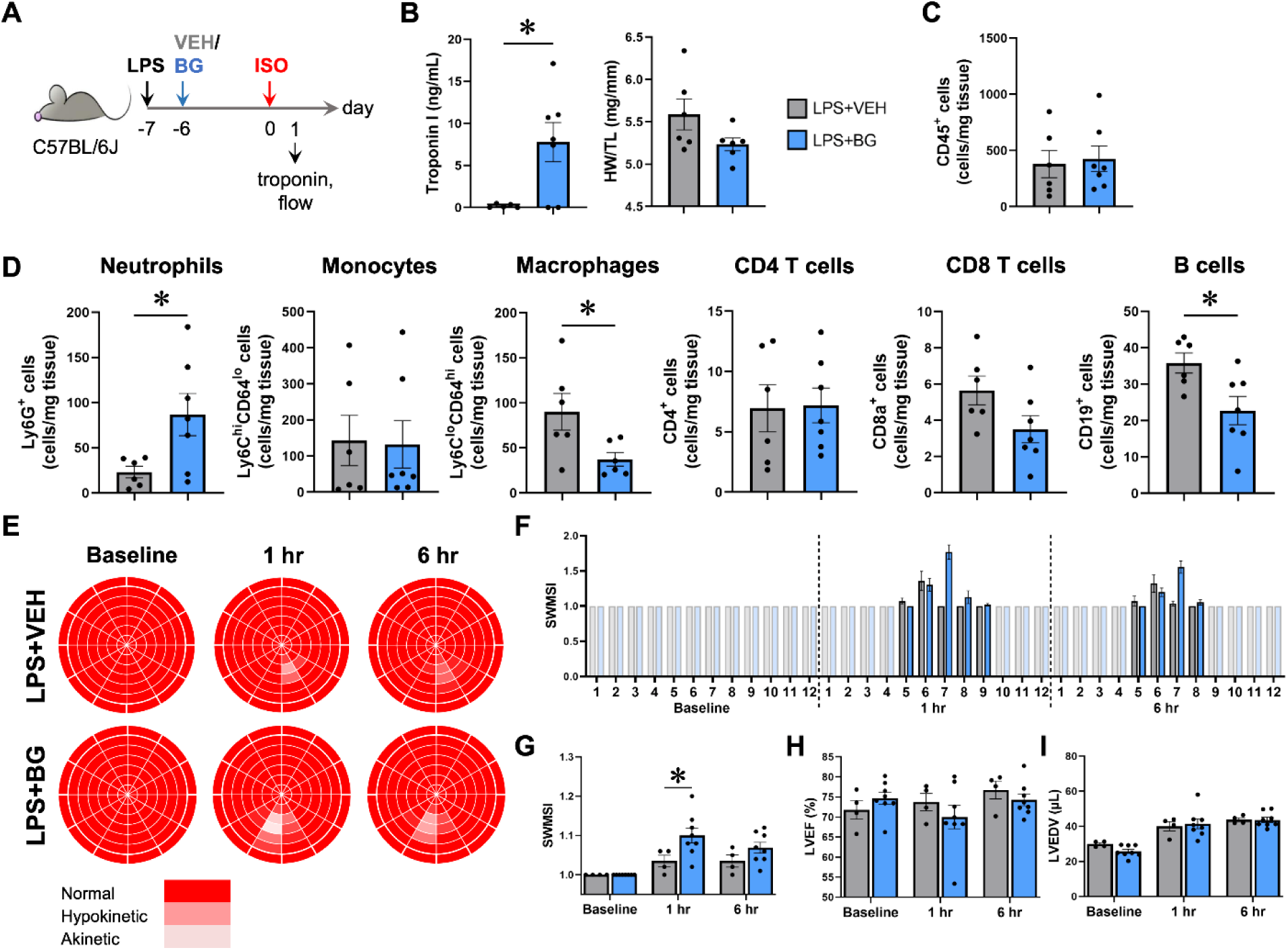
Effects of β-glucan treatment on LPS-induced cardioprotection. **(A)** Mice were injected i.p. with lipopolysaccharide (LPS; 2.5 mg/kg) on day -7, β-glucan (BG; 1 mg/mouse) or vehicle (VEH) on day -6, and then challenged with ISO (300 mg/kg) on day 0. All mice were evaluated on day 1. **(B)** Serum cardiac troponin I levels and heart weight-to-tibia length ratios in LPS+VEH and LPS+BG mice. Flow cytometric analysis of **(C)** total CD45^+^ cells, **(D)** Ly6G^+^ neutrophils, Ly6C^hi^CD64^lo^ monocytes, Ly6C^lo^CD64^hi^ macrophages, CD4^+^ T cells, CD8a^+^ T cells, and CD19^+^ B cells in the heart on day 1, with counts expressed as cells/mg of tissue (n=6-7/group). Cardiac function was also evaluated in LPS+VEH and LPS+BG mice using 2-D echocardiography. **(E)** Bulls eye regional wall motion plots of the left ventricle. **(F)** Quantification of the segmental wall motion score index (SWSMI) per segment in (E). **(G)** The global SWMSI was determined as the average of 84 regional LV segments, where 1 = normal wall thickening, 2 = hypokinesis, 3 = akinesis **(H)** Left ventricular ejection fraction (LVEF), and **(I)** left ventricular end-diastolic volume (LVEDV) measurements (n=4-8/group). Data show mean ± SEM and were analyzed by unpaired, two-tailed t-test (B-D), or repeated measures two-way ANOVA with Sidak’s multiple comparisons test (G-I). (Key: \**P*<0.05).

### Multiomic analysis of LPS-induced tolerance

To identify putative cell types and genes that contributed to LPS-induced tolerance against ISO-injury, we performed multiome single nuclei ATAC-seq (snATAC-seq) and single nuclei RNA-seq (snRNA-seq) on three groups of mice: mice treated with LPS on day -7 and vehicle on day -6 (tolerized), mice treated with LPS on day -7 and β-glucan on day -6 (reversal of tolerance), or mice treated with PBS on day -7 and vehicle on day -6 (control). All hearts were harvested on day 0 for nuclei isolation and sequencing (**Figure 5A**). **Figure 5B** shows a UMAP plot of the snRNA-seq data, whereas **Figure 5C** summarizes the frequencies of the various cell types in the control, tolerized, and reverse-tolerized groups of mice. As shown, there were no obvious differences in the frequencies of the different cell types in the control, tolerized and reverse-tolerized mouse hearts consistent with our flow cytometry data. Next, we compared the number of differentially expressed genes (DEGs) in the 7 predominant cell types in tolerized hearts compared to hearts with reversal of LPS tolerance. As shown in **Figure 5D**, there was a substantial number of DEGs in the cell groups from tolerized mouse hearts. Moreover, all cell groups in the tolerized mouse hearts exhibited increased > decreased gene expression. The number of total DEGs was greatest for endothelial cells > endocardial cells > myeloid cells > fibroblasts > pericytes > cardiac myocytes > T and Natural Killer (TNK) cells. Next, we interrogated the snATAC-seq data to determine whether there were differences in the number of peaks with increased chromatin accessibility across the different cell types. As shown in **Figure 5E**, the tolerized state was characterized by increased numbers of accessible chromatin peaks across all cell types. Of note the degree of change in chromatin peak accessibility was cell-type dependent, with increased accessibility observed in TNK cells > endothelial cells > myeloid cells > pericytes > cardiac myocytes > endocardial cells > fibroblasts. Heatmaps of differentially accessible chromatin peaks revealed that the cells from tolerized hearts had increased numbers of accessible chromatin peaks relative to control hearts, whereas cells from hearts that had reversed tolerance were distinct from those in tolerized hearts and were more similar to control hearts **(Figure 5F)**. Next, we leveraged the RNA/ATAC information from the same nucleus to link these regulatory elements to putative target genes. Using the genes linked to differential regulatory elements, we performed a pathway analysis (**Supplemental Table S1**) in order to evaluate the different biological themes that characterized the various cell types in tolerized mouse hearts. As shown in **Figures 5G-5M**, the different cell types in the heart exhibited distinct sets of upregulated biological pathways. However, the important finding illustrated in **Figures 5G-5M** is that pathways related to an interferon response were overrepresented across all cell types in the heart, suggesting that increased interferon signaling encodes key epigenetic changes and is a biological theme that is common to cell types under LPS-induced tolerance to ISO-injury.

**Figure 5.**
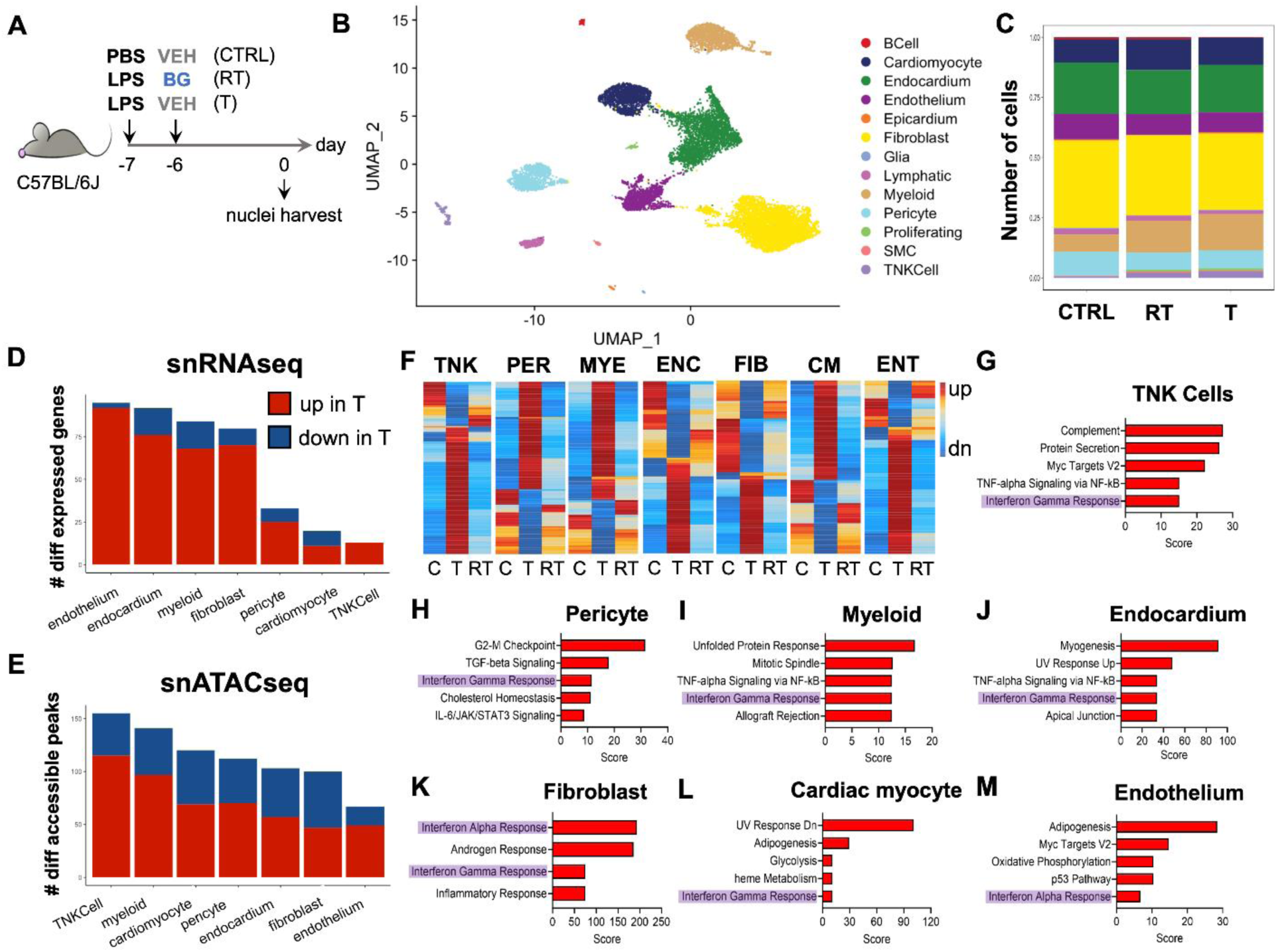
Multiomic analysis of LPS-induced cardioprotection. **(A)** Mice were injected i.p. with saline (PBS) on day -7 and vehicle (VEH) on day -6 (control, CTRL), lipopolysaccharide (LPS; 2.5 mg/kg) on day -7 and β-glucan (BG; 1 mg/mouse) on day -6 (reverse-tolerized, RT), or LPS on day -7 and VEH on day -6 (tolerized, T), and nuclei were harvested from hearts on day 0 for multiomic snRNA-seq and snATAC-seq analysis (nuclei pooled from n=3/group). **(B)** UMAP plot showing the total cell populations found in CTRL, T, and RT hearts. **(C)** Quantified cell composition from (B). **(D)** Number of differentially expressed genes in the tolerized group, compared to the reverse-tolerized group. **(E)** Number of differentially accessible chromatin peaks in the tolerized mice, compared to the reverse tolerized mice. **(F)** Heat maps of differentially accessible peaks across TNK cells (TNK) pericytes (PER), myeloid cells (MYE), endocardial cells (ENC), fibroblasts (FIB), cardiac myocytes (CM), and endothelial cells (ENT) in the CTRL (C), tolerized and reverse-tolerized groups. **(G-M)** Pathways enriched from genes linked to chromatin peaks with increased accessibility in the tolerized mouse hearts compared with the reverse tolerized mouse hearts group for the 7 predominant cell types in the heart.

To confirm the findings with respect to increased expression of interferon-related signaling genes, we performed RT-qPCR in LPS-treated hearts on select candidate genes that either initiated or were downstream of type 1 interferon signaling (*Ifnb1* and *Ifna2, Ifit1),* type 2 interferon signaling (*Ifng)*, and type 1/2 interferon signaling (*Irf1*). As shown in **Supplemental Figure S4**, we confirmed that there was increased expression of the type 1 interferon-specific downstream target gene *Ifit1* (*P*=0.006) and the type 1/2 interferon downstream target gene *Irf1* (*P*=0.0008) on day 1 in LPS-treated mice.

### Type 1/Type 2 interferon signaling is required for the cardioprotective effects of LPS

Based upon our bioinformatics analysis which suggested that genes related to interferon signaling pathways were enriched in all cell types in the LPS-tolerized mouse hearts (**Figure 5G-5M**), as well as prior studies which suggested that interferon-induced signaling increases the expression of genes involved in cellular protection and survival,^20^ we asked whether type 1 or type 2 interferon, or combined type 1/type 2 interferon signaling was required for LPS-induced tolerance against ISO-injury. As shown in **Figure 6A**, mice were first pretreated with LPS on day -7 followed by treatment 1 hour later with monoclonal Abs (mABs) against the type 1 interferon receptor (IFNAR1), the type 2 interferon receptor (IFNγR), or both interferon receptors (IFNAR1/IFNγR). IgG1 (for anti-IFNAR1), IgG2a (for anti-IFNγR) mABs and IgG1/IgG2a mAbs were used as the appropriate isotype control antibodies. Mice were subjected to ISO-injury on day 0 and then phenotyped on day 1. As shown in **Supplemental Figure S5** blocking either the type 1 or the type 2 interferon receptor using concentrations of mABs that were sufficient to downregulate type 1/2 interferon signaling did not attenuate LPS-induced tolerance against ISO-injury. In contrast, blocking both the type 1 and type 2 interferon receptors using identical concentrations of mABs (**Figure 6B**) resulted in a significant increase (*P*=0.014) in serum cardiac troponin I levels on day 1 after ISO-injection (**Figure 6C**), as well as a significant increase (*P*=0.002) in infiltrating neutrophils on day 1 (**Figure 6E**), suggesting attenuation of LPS-induced tolerance against ISO-injury. The finding that blocking both the type 1 and type 2 interferon receptors was required to block LPS-induced tolerance is consistent with the observation that there is interaction (i.e. crosstalk) between the type 1 and type 2 interferon receptor downstream signaling pathways.^21^

**Figure 6.**
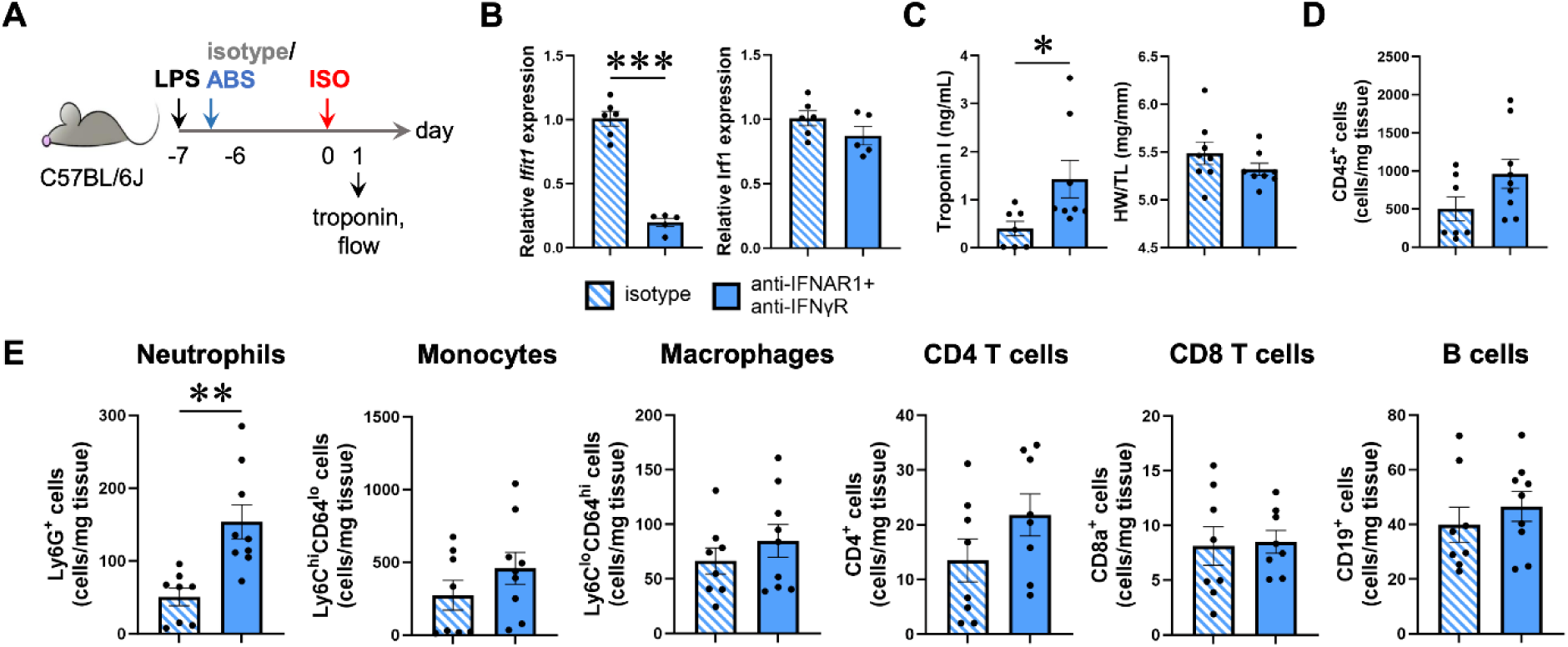
Effects of blocking interferon receptor signaling on LPS-induced cardioprotection. **(A)** Mice were injected i.p. with lipopolysaccharide (LPS; 2.5 mg/kg) on day -7, antibodies blocking the interferon receptors IFNAR-1 and IFNγR (ABS) or their corresponding isotype controls (500 μg/mouse) an hour after the LPS dose, and then challenged with ISO (300 mg/kg) on day 0. All mice were evaluated on day 1. **(B)** *Ifit1* and *Irf1* gene expression as evaluated by qPCR in isotype and blocking antibody-treated mice (n=5-6/group). **(C)** Serum cardiac troponin I levels and heart weight-to-tibia length ratios. Flow cytometry analysis of **(D)** total CD45^+^ cells, **(E)** Ly6G^+^ neutrophils, Ly6C^hi^CD64^lo^ monocytes, Ly6C^lo^CD64^hi^ macrophages, CD4^+^ T cells, CD8a^+^ T cells, and CD19^+^ B cells in the heart on day 1 (n=8-9/group). Data show mean ± SEM and were analyzed by unpaired, two-tailed t-test (B, C, for HW/TL, D-E), or the Mann-Whitney test (C, for troponin). (Key: \**P*<0.05, \*\**P*<0.01, \*\*\**P*<0.001)

To complement the above loss-of-function studies, we performed gain of function studies by pre-treating mice with recombinant murine interferon-β1 (type 1 interferon), recombinant murine interferon-γ (type 2 interferon) or a combination of recombinant murine interferon-β1/interferon-γ 7 days prior to ISO-injection (**Figure 7A)**; diluent (PBS) treated mice served as the appropriate controls. As expected, pre-treatment with recombinant interferons increased the expression of *Ifit1* in the heart (**Figure 7B**). Consistent with above loss-of-function studies, pre-treating the mice with recombinant murine interferon-β1 or interferon-γ did not protect against the ISO-induced increases in cardiac troponin I levels when compared to diluent-treated control mice (*P*=0.157 and *P*=0.196, respectively) (**Figure 7C**). Similarly, pre-treating the mice with interferon-β1 or interferon-γ alone did not reduce the number of total CD45^+^ cells (*P*=0.992 and *P*=0.965, respectively) and infiltrating Ly6G^+^ neutrophils (*P*=0.960 and *P*=0.901, respectively) in the heart after ISO-injection when compared to diluent-treated control mice (**Figure 7D-7E**). In contrast, we observed a significant decrease in troponin I levels (*P*=0.007) on day 1 in the ISO-injected mice that had been pre-treated with a combination of interferon-β1 and interferon-γ (**Figure 7C**), as well as significant decreases in the number of total CD45^+^ cells (*P*=0.020) and infiltrating neutrophils (*P*=0.005) (**Figure 7D-7E**). There were no other significant changes observed in the other immune cell subsets in the heart in any of the experimental groups that were studied (**Figure 7E**).

**Figure 7.**
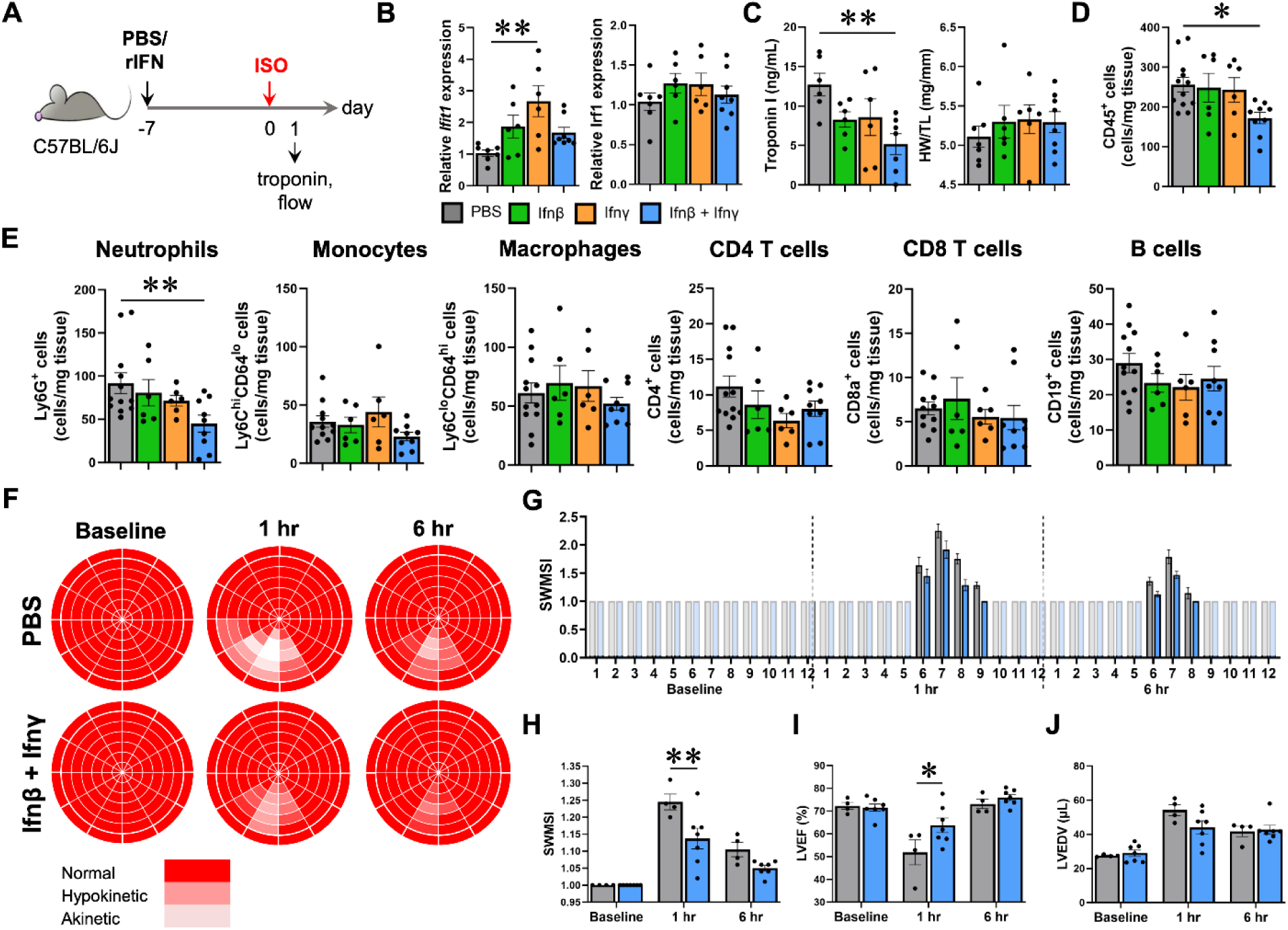
Recombinant interferon treatment induces cardioprotection. **(A)** Mice were injected i.p. with recombinant murine interferon-β1 (1 μg), -γ (10 μg), both, or saline (PBS) on day -7 and then challenged with ISO (300 mg/kg) on day 0. All mice were evaluated on day 1. **(B)** *Ifit1* and *Irf1* gene expression as evaluated by qPCR across groups. **(C)** Serum cardiac troponin I levels and heart weight-to-tibia length ratios. Flow cytometry analysis of **(D)** total CD45^+^ cells, **(E)** Ly6G^+^ neutrophils, Ly6C^hi^CD64^lo^ monocytes, Ly6C^lo^CD64^hi^ macrophages, CD4^+^ T cells, CD8a^+^ T cells, and CD19^+^ B cells in the heart on day 1 (n=6-8/group). Cardiac function was also evaluated in mice injected with both interferons β1 and γ and compared with PBS-injected mice using 2-D echocardiography. **(F)** Bulls eye regional wall motion plots of the left ventricle. **(G)** Quantification of the segmental wall motion score index (SWSMI) per segment in (E). **(H)** The global SWMSI was determined as the average of 84 regional LV segments, where 1 = normal wall thickening, 2 = hypokinesis, 3 = akinesis. **(I)** Left ventricular ejection fraction (LVEF) and **(J)** left ventricular end-diastolic volume (LVEDV) (n=4-7/group). Data show mean ± SEM and were analyzed by the Kruskal-Wallis test with Dunn’s multiple comparisons test (B, for *Ifit1*), the one-way ANOVA with Dunnett’s multiple comparisons test (B, for *Irf1*, C-E) or repeated measures two-way ANOVA with Sidak’s multiple comparisons test (H-J). (Key: \**P*<0.05, \*\**P*<0.01)

To determine if treatment with recombinant interferon was sufficient to attenuate the LPS-induced cardioprotective effects on LV structure and function, we performed 2-D echocardiography at baseline and at 1 and 6 hours after ISO injection. As shown in **Figures 7F-7H** there was a decrease in the extent of ISO-induced segmental wall motion abnormalities in the mid to apical inferior LV segments and a significant decrease (*P*=0.002) in the SWMSI score at 1 hour in the interferon-β1/interferon-γ treated mice relative to diluent control mice. There was also a significant increase in LVEF (*P*=0.015) at 1 hour in the interferon-β1/interferon-γ treated mice relative to diluent control mice **(Figure 7I)**, whereas there was no significant difference (*P*=0.443, by two-way ANOVA) in the LVEDV between diluent treated control and interferon-β1/interferon-γ treated mice **(Figure 7J)**. Viewed together these results suggest that LPS-induced tolerance against ISO-mediated cardiac injury is mediated, at least in part, through increased interferon type1/type 2 signaling.

### Multiomic analysis of interferon-induced cardioprotection

To identify the cell types and genes contributing to IFN-induced tolerance against ISO injury, we performed a multiomic snATAC-seq and snRNA-seq analysis of mice treated with recombinant murine interferon-β1/γ on day -7 **(Supplemental Figure S6A**) and mice treated with diluent on day -7 (data from **Figure 5**). We observed differentially accessible chromatin peaks in all cell types examined (**Supplemental Figure S6B**). The peaks with increased accessibility varied across cell types and was greatest for cardiac myocytes > TNK cells > pericytes, myeloid cells, fibroblasts > endothelial cells > endocardial cells (**Supplemental Figure S6B**), which was different from what we observed in LPS-tolerized hearts **(Figure 5)**. Of interest, the chromatin accessibility heat maps in the different cell populations from IFN-treated hearts displayed a pattern that was the inverse of the chromatin accessibility landscapes observed in cell groups from the control hearts (**Supplemental Figure S6C).** We performed a pathway analysis of genes associated with chromatin peaks exhibiting increased accessibility in the major cell types from the IFN-treated mouse hearts (**Supplemental Table S2**), to explore the biological themes which characterized the different cell types. The pathways that were identified were distinct in each of the different groups of cells (**Supplemental Figure S6D**), with Hedgehog signaling noted as a prominent biological theme in cardiac myocytes, pericytes, and myeloid cells, and IFN-mediated signaling noted in myeloid cells and endothelial cells. Given that Hedgehog signaling has been implicated in maintaining tissue homeostasis after cardiac injury,^22, 23^ we examined the DEGs that were associated with the Glioma-associated oncogene homologue 1 (Gli1), Gli2, and Gli3 family of transcription factors, which are the primary transcription factors involved in Hedgehog signaling (**Supplemental Table S3**). As depicted in the heat map in **Supplemental Figure S6E,** there were increased DEGs in all cell types, with the greatest number of DEGs in cardiac myocytes > endocardial cells > fibroblasts > myeloid cells, consistent with our bioinformatic analysis of actively transcribed genes in chromatin regions with increased accessibility **(Supplemental Figure S6D)**. Lastly, we examined DEGs in the cell types in IFN-treated hearts that were involved in the biological process ontologies of cellular response to stress (GO:0080135) and wound healing (GO:0042060) (**Supplemental Figures S6F-S6G)**. Remarkably, the cell types with the greatest number of DEGs in these two gene ontology categories coincided with the cell types that had the greatest number of DEGs linked to the Gli transcription factors downstream of Hedgehog signaling.

### LPS-induced cardioprotection is associated with changes in intercellular communication networks

The multiomic snRNA-seq and snATAC-seq data (**Figures 5D-E**) in LPS-tolerized hearts revealed differences in gene expression and chromatin peak accessibility across all cell types, with the most significant changes observed in non-cardiac myocyte cells. These findings suggested a potential model wherein LPS induces cardioprotection through coordinated paracrine responses that are provided by non-myocyte cell types in the heart. To begin to explore this possibility, we used unsupervised CellChat^11^ to analyze intercellular communication networks in the LPS-tolerized hearts, as well as hearts that had reversal of LPS tolerance. Intriguingly, we discovered that both the number and the strength of cellular interactions (**Figure 8A**) between cardiac myocytes and other cell types were increased, especially with TNK cells, myeloid cells, pericytes, and fibroblasts (**Figure 8B**). We also noted increased autocrine signaling in cardiac myocytes; however, the strength of the autocrine signaling was less than what was observed with paracrine signaling from non-myocyte cell types. A second interesting finding was that both the number and strength of cell-cell interactions between fibroblasts and other cell types in the heart were decreased in the LPS-tolerized hearts when compared to hearts with reversal of LPS tolerance.

**Figure 8.**
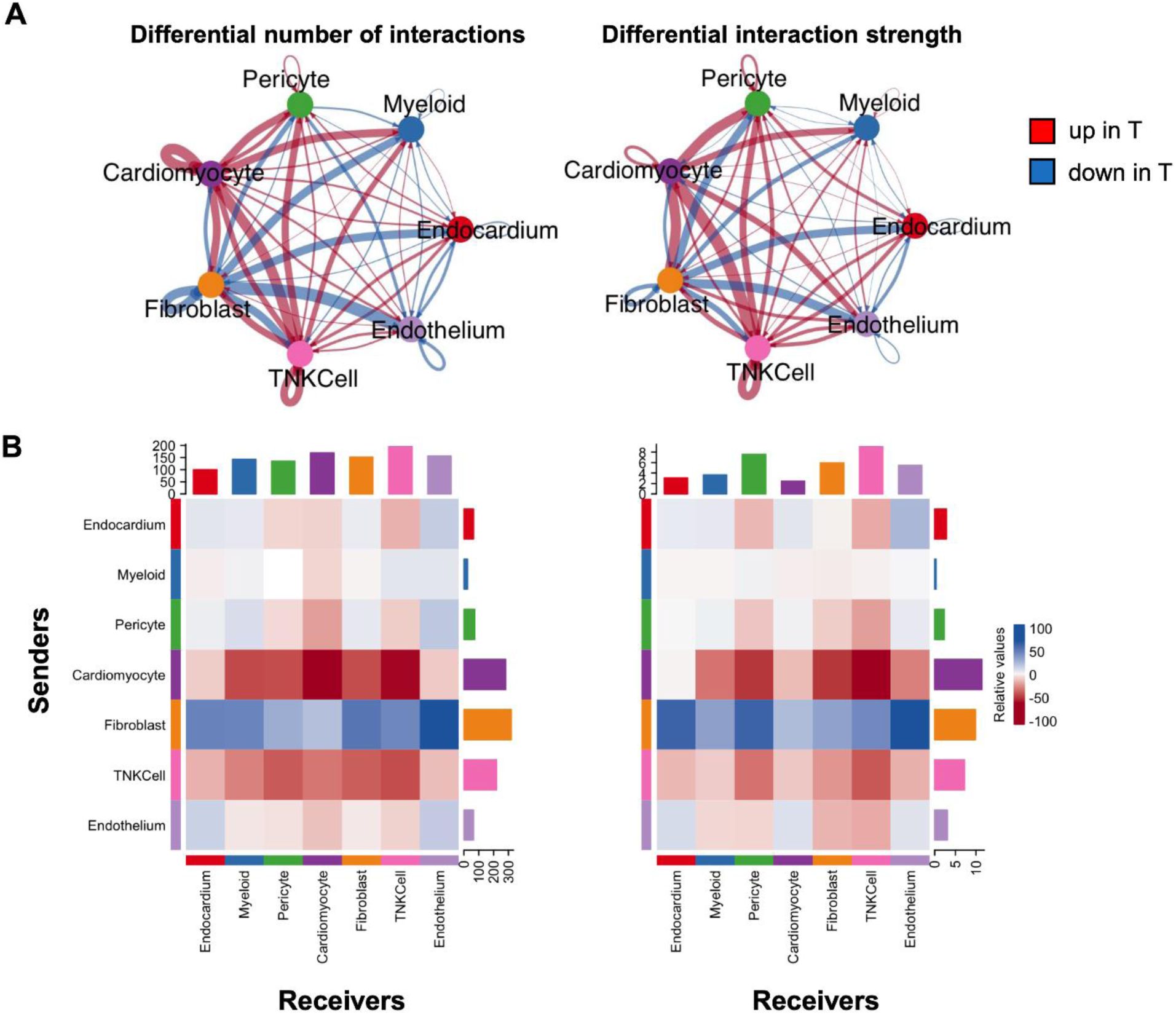
Analysis of cell-cell communication networks in LPS-tolerized hearts. **(A)** CellChat was used on the multiomic data generated in Figure 5 to create cell-cell communication network maps between the 7 predominant cell types in the heart, comparing LPS-tolerized mice to those with treated with LPS + β-glucan. **(B)** Heat map visualization of the communication networks in (A), with sender cells on the y-axis and receiver cells on the x-axis. For (A) and (B), left panels indicate the number of interactions, whereas right panels indicate the strength of interactions.

## Discussion

The findings of this study demonstrate that pretreatment with LPS cross-tolerizes the heart against the myopathic effects of ISO through epigenetic regulation of IFN type 1/type 2-mediated signaling. The following lines of evidence support this statement. First, pretreatment with LPS prevented the myopathic effects of ISO, including decreased ISO-induced cardiac myocyte cell death **(Figure 2)**, reduced inflammation **(Figure 2)**, preserved LV structure and function **(Figure 3)** and increased survival **(Figure 2)**. Second, the protective effects of LPS were reversed by β-glucan, as evidenced by the ISO-induced increase in troponin release, increase in Ly6G^+^ neutrophil infiltration, as well as an increase in ISO-induced LV regional wall motion abnormalities **(Figure 4)**. Third, multiomic analysis of hearts treated with LPS and LPS+β-glucan revealed that the LPS-induced tolerance against ISO injury was encoded by epigenetic changes at the chromatin level in immune cells, stromal cells, and myocytes. Linking gene regulatory elements to putative genes revealed that the interferon signaling pathway was enriched in all cell types in LPS-tolerized mouse hearts **(Figure 5)**, in line with studies showing that TLR4-mediated signaling through the MyD88-independent TRIF-dependent pathway results in enhanced interferon signaling.^4^ Remarkably, both loss- **(Figure 6)** and gain- **(Figure 7)** -of-function studies demonstrated that interferon type 1/type 2 signaling was necessary and sufficient to mediate the cytoprotective effects of LPS, consistent with studies that have shown that type 1^24^ and type 2 interferon signaling is cardioprotective.^25^ Fourth, multiomic analysis of interferon type 1/2-treated mouse hearts was remarkable for showing increased chromatin accessibility in cardiac myocytes, fibroblasts and endothelial cells, as well as an upregulation of genes involved in cardiac repair (e.g., Hedgehog signaling, Notch signaling) **(Supplemental Figure S6)**. Finally, computational analysis of cell-cell communication networks in the LPS-treated hearts was notable for increased paracrine signaling in cardiac myocytes and cardiac stromal cells, especially with TNK cells, myeloid cells, pericytes, and fibroblasts **(Figure 8)**. Although autocrine signaling was increased in cardiac myocytes and TNK cells, the strength of the autocrine signaling was comparatively less than the strength of paracrine signaling in cardiac stromal cells. Viewed together these results suggest a hypothetical model of LPS-induced cross-tolerance, wherein LPS provokes chromatin remodeling and increased interferon type 1/type 2 autocrine/paracrine signaling in cardiac myocytes and stromal cells. These changes result in the upregulation of cytoprotective gene networks that enhance the resilience of cardiac myocytes against external stressors (e.g., neurohormonal stress).

### Trained Innate Immune Tolerance in the Heart

In a seminal study Murry, Jennings, and Reimer demonstrated that brief episodes of repetitive ischemia protected the heart against a subsequent bout of prolonged ischemia, which was referred to as “preconditioning.”^26^ Ensuing studies showed that, in addition to ischemic injury, various other stimuli were able to protect the heart against subsequent episodes of prolonged ischemia (reviewed in ^27^ ^28^). A closer examination of the mechanisms of action of preconditioning suggests that this phenomenon shares a number of overlapping biological themes with trained innate immune responses.^29^ Trained innate immunity refers to the long-lasting functional reprogramming of innate immune cells triggered by an external or internal stressor. This reprogramming alters the response of the innate immune system to a subsequent second challenge, either amplifying the response (trained immunity) or dampening the response (immune tolerance). The functional reprogramming in cells is driven by epigenetic, transcriptional and metabolic reprogramming,^29^ which can be induced by exogenous microbial ligands (e.g., PAMPs), or by endogenous ligands (e.g. DAMPS) that are recognized by pattern recognition receptors, such as TLRs, C-type lectin receptors, and NOD-like receptors. Cytokines that are released during an immune response can also contribute to trained immunity by affecting the programming of immune cells in an autocrine or paracrine manner.^30^

The physiological role of innate immune tolerance is to limit the extent of tissue damage that occurs in the setting of prolonged or excessive inflammation. One of the hallmarks of innate immune tolerance is its lack of specificity, insofar as exposure to one ligand can induce a state of protection against a different ligand (referred to as cross-tolerance).^31^ Intriguingly, the induction of innate immune memory is not limited to immune cells but can also occur in stromal and epithelial cells, which maintain open chromatin in key stress response genes that are activated by the primary stimulus.^32, 33^ Here we show that trained innate immune responses also occur in cardiac myocytes. The tolerance that develops in trained innate immunity is primarily cell-autonomous, based on studies demonstrating that the epigenetic changes that occur within immune cells, such as histone methylation and acetylation, do not require external signals from other cell types to be maintained, as well as the observation that epigenetic modifications can be passed onto the progeny of cells.^34^ While innate immune tolerance is primarily cell-autonomous, there are likely non-cell-autonomous mechanisms that may also contribute to the development of tolerance, including the secretion of cytokines (e.g., interferons^35^) or other soluble factors that can modulate the function of neighboring cells, thereby contributing to a broader immune response within the tissue microenvironment.^34^ The results of the present study suggest that LPS induced tolerance in the heart is mediated by cell-autonomous and non-cell-autonomous mechanisms.

The findings in this study are consistent with our prior observation that ISO-induced tissue injury protects the heart from the myopathic effects of a second exposure to ISO.^2^ The potential common mechanistic link between our past and current work is that LPS is a classic TLR4 ligand, whereas ISO-mediated cardiac myocyte necrosis leads to the release of endogenous DAMPs that also signal through TLR4. Given that we used the standard preparation of LPS, we cannot exclude that some of the observed effects were non-TLR4 effects. Our studies are also in overall agreement with prior studies showing that pretreatment with LPS 24 hours prior to ischemic injury protected the heart against ischemia/reperfusion injury when the hearts were evaluated immediately after ischemic tissue injury.^36–40^ However, when LPS was administered 1-2 hours prior to ischemic injury, the cytoprotective effects of LPS were not evident in vivo,^36, 39^ which gave rise to the concept that LPS induced late preconditioning in the heart. Our studies both confirm and expand upon these prior studies by demonstrating that LPS pretreatment 7 days prior to cardiac injury induces chromatin remodeling and upregulation of the expression of interferon response genes in cardiac myocytes and stromal cells in the heart. Additionally, we show that type 1 and type 2 interferon signaling is both necessary and sufficient for the late cytoprotective effects of LPS, which is in line with prior studies that have implicated a cytoprotective role for activation of the Janus kinase (JAK) and the signal transducer and activator of transcription (STAT) in the setting of ischemia/reperfusion injury (i.e., SAFE pathway).^41, 42^

## Conclusion

This study shows that pretreatment with LPS induces cross-tolerance against the cytotoxic effects of ISO in the heart, consistent with trained innate immune tolerance in the heart. While efforts to therapeutically leverage trained immune responses have traditionally focused on heightened immune responses (e.g. vaccine development), there is growing interest in exploring the role of trained immune tolerance as a therapeutic approach to treating allergic reactions, autoimmune disease, and transplant organ survival.^43^ However, as illustrated by our observation that the cytoprotective effects of LPS in cardiac myocytes are both cell-autonomous and non-cell autonomous, it is likely that the signaling pathways that are involved in the development of innate immune tolerance are highly complex and not fully understood. Going forward it will be important to understand what constitutes a tolerance phenotype (i.e., functional state) in innate immune cells, insofar as the molecular mechanisms underlying tolerance (e.g., epigenetic reprogramming) likely overlap those of non-tolerized innate immune cells. Thus, deciphering the cell signatures that define tolerance will be essential for developing targeted approaches to reprogram, regulate, or selectively ablate immune cells to restore tissue homeostasis following tissue injury. Although speculative, one possible explanation for why preconditioning studies have not (yet) translated into improved outcomes in clinical trials^44,45^ is that they have focused more on the requisite nature of the preconditioning stimulus, rather than focusing on the molecular signatures of the tolerance phenotype(s) in the heart. Further studies will be necessary to address these interesting, if not important questions.

## Acknowledgments

We would like to thank the Genome Technology Access Center at the McDonnell Genome Institute at Washington University School of Medicine for help with genomic analysis. The Center is partially supported by NCI Cancer Center Support Grant #P30 CA91842 to the Siteman Cancer Center.

