## Supplemental Data for "Lipopolysaccharide Induces Trained Innate Immune Tolerance in the Heart Through Interferon Signaling in a Model of Stress-Induced Cardiomyopathy"

### Table of Contents for Supplemental Appendix


| Supplemental Appendix | Page |
| --- | --- |
| **Supplemental Figure S1.** Gating strategy for flow cytometry counting of immune cells……………………………………………………………………………………. | 2 |
| **Supplemental Figure S2.** Comparison of cardioprotection induced by crude versus ultrapure lipopolysaccharide……………………………………………………………... | 3 |
| **Supplemental Figure S3.** Characterization of immune cells in the blood and spleen in LPS pretreated mice following ISO injury…………………………………………….. | 4 |
| **Supplemental Figure S4.** Quantification of interferon-related gene expression in LPS pretreated hearts…………………………………………………………………………….. | 5 |
| **Supplemental Figure S5.** Effects of blocking individual interferon receptor signaling on LPS-induced cardioprotection………………………………………………………... | 6 |
| **Supplemental Figure S6.** Multiomic analysis of recombinant interferon-treated hearts. | 7 |

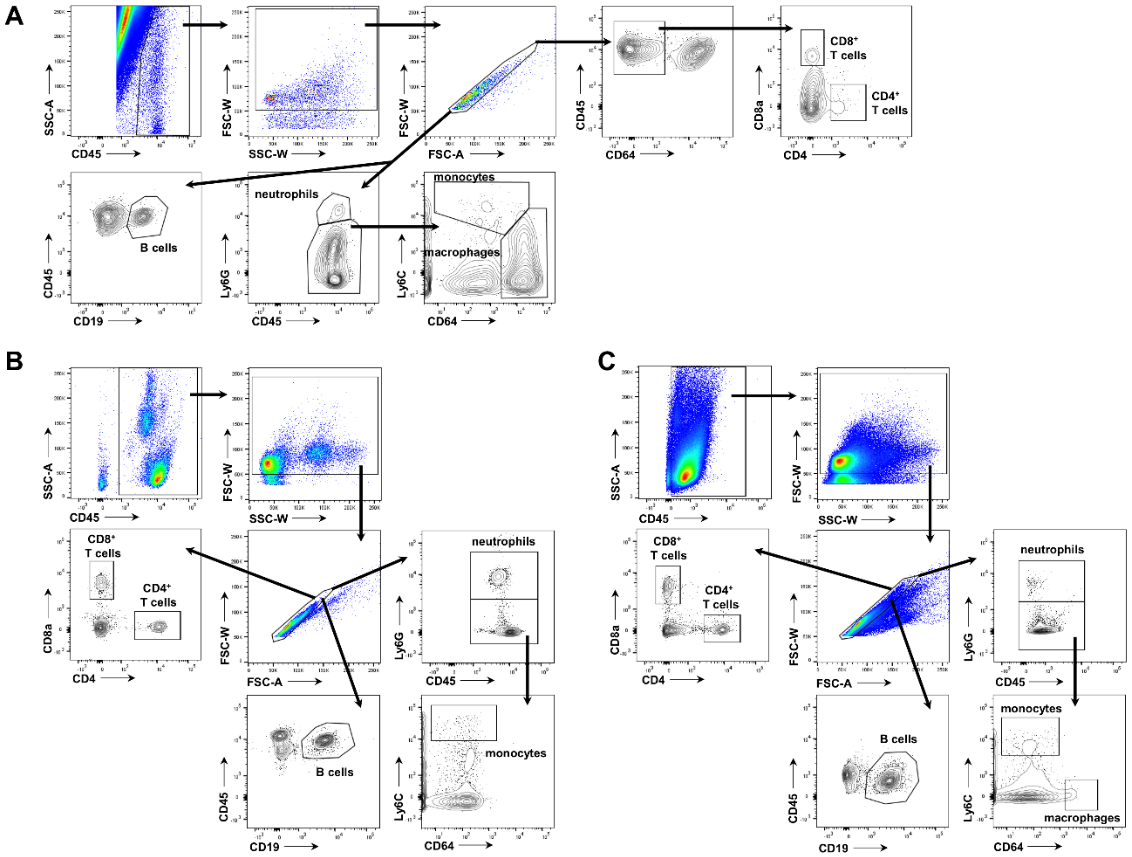


**Supplemental Figure S1. Gating strategy for flow cytometry counting of immune cells.** Representative gating strategy for enumerating CD45$^{+}$ cells, Ly6G$^{+}$ neutrophils, Ly6C$^{hi}$CD64$^{lo}$ monocytes, Ly6C$^{lo}$CD64$^{hi}$ macrophages, CD4$^{+}$ T cells, CD8$^{+}$ T cells, and CD19$^{+}$ B cells in the **(A)** heart, **(B)** blood, and **(C)** spleen.

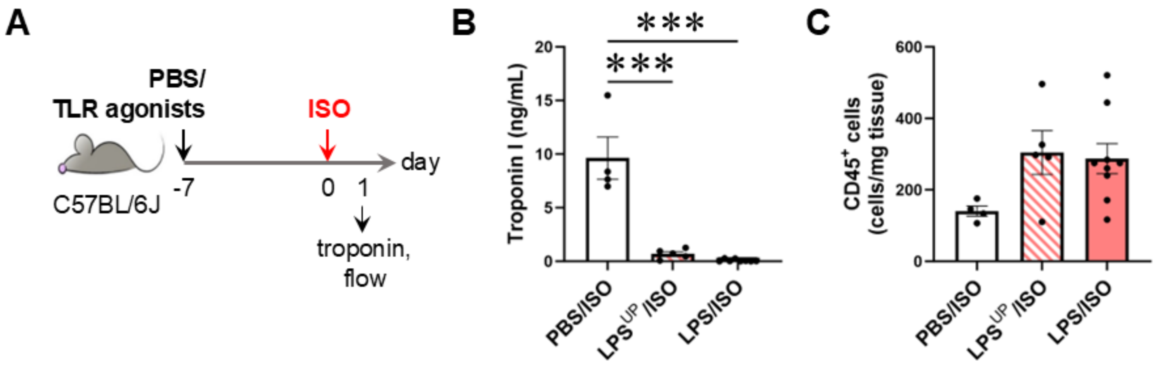


**Supplemental Figure S2. Comparison of cardioprotection induced by crude versus ultrapure lipopolysaccharide. (A)** Mice were injected i.p. with crude lipopolysaccharide (LPS) or ultrapure LPS (LPS$^{UP}$), both at 2.5 mg/kg, or saline (PBS) on day -7 and challenged with an i.p. injection of ISO (300 mg/kg) on day 0. Mice were then evaluated on day 1 post-ISO. **(B)** Serum cardiac troponin I analysis. **(C)** Flow cytometry analysis of total CD45$^{+}$ cells in the heart on day 1 (n=4-9/group). Data show mean ± SEM and were analyzed by one-way ANOVA with Tukey's post-hoc test (**B-C**). Data for the LPS/ISO group were from the same mice used in **Figure 1**. (Key: ****P*<0.001)

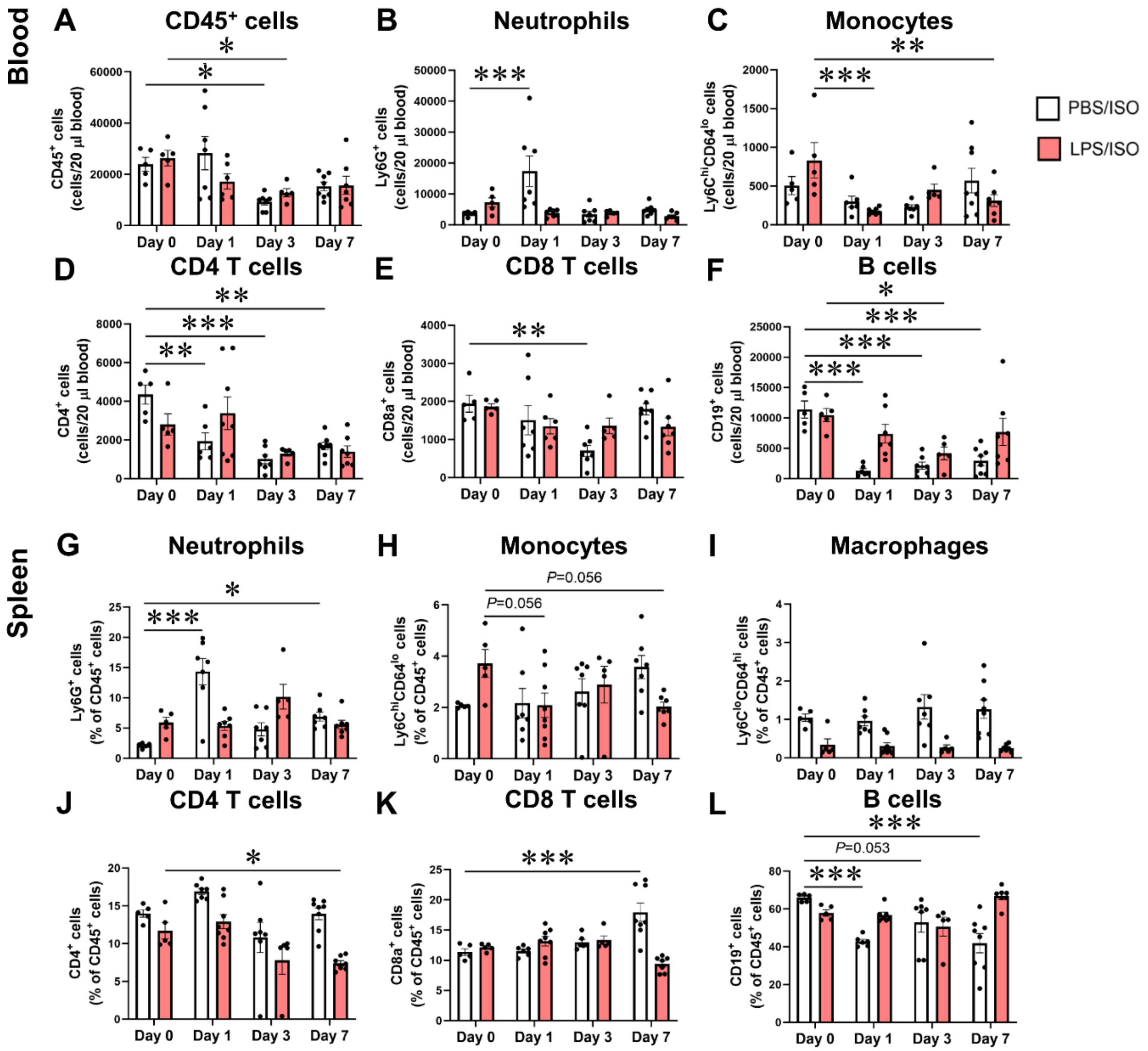


**Supplemental Figure S3. Characterization of immune cells in the blood and spleen in LPS pretreated mice following ISO injury.** Mice were injected i.p. with lipopolysaccharide (LPS; 2.5 mg/kg) or saline (PBS) on day -7 and challenged with an i.p. injection of ISO (300 mg/kg) on day 0. PBS/ISO and LPS/ISO mice were evaluated on day 1. Flow cytometry analysis of **(A)** total CD45$^+$ cells, **(B)** Ly6G$^+$ neutrophils, **(C)** Ly6C$^{hi}$CD64$^{lo}$ monocytes, **(D)** CD4$^+$ T cells, **(E)** CD8a$^+$ T cells, and **(F)** CD19$^+$ B cells in the blood, and of **(G)** Ly6G$^+$ neutrophils, **(H)** Ly6C$^{hi}$CD64$^{lo}$ monocytes, **(I)** Ly6C$^{lo}$CD64$^{hi}$ macrophages, **(J)** CD4$^+$ T cells, **(K)** CD8a$^+$ T cells, and **(L)** CD19$^+$ B cells in the spleen (n=5-8/group). Data show mean ± SEM and were analyzed by two-way ANOVA with Dunnett's multiple comparisons test. (Key: **P*<0.05, ***P*<0.01, ****P*<0.001)

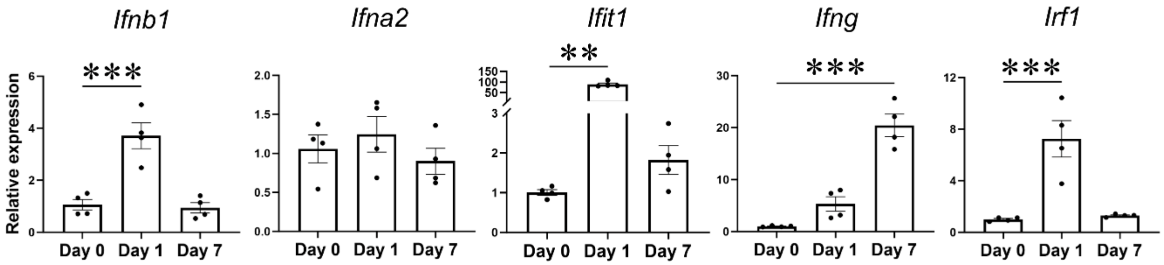


**Supplemental Figure S4. Quantification of interferon-related gene expression in LPS pretreated hearts.** Mice were injected i.p. with lipopolysaccharide (LPS; 2.5 mg/kg) on day 0. Hearts were collected at baseline (day 0), and at days 1 and 7 post-injection, and used for qPCR analysis of *Ifnb1*, *Ifna2*, *Ifit1*, *Ifng*, and *Irf1* (n=4/group). Expression was normalized to *Gapdh* and calculated relative to baseline. Data show mean ± SEM and were analyzed by one-way ANOVA with Dunnett's multiple comparisons test for all genes, except for *Ifit1*, which was analyzed by the Kruskal-Wallis test with Dunn's multiple comparisons test. (Key: $**P<0.01$, $***P<0.001$)

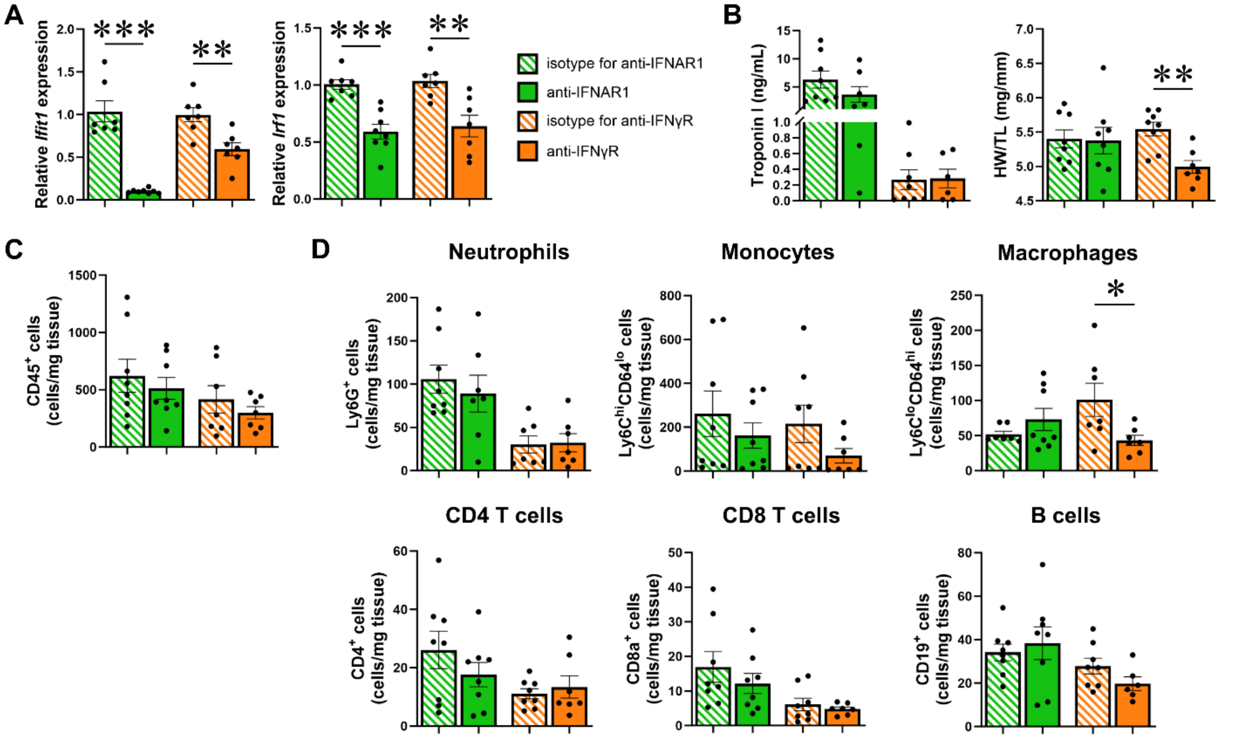


**Supplemental Figure S5. Effects of blocking individual interferon receptor signaling on LPS-induced cardioprotection.** Mice were injected i.p. with lipopolysaccharide (LPS; 2.5 mg/kg) on day -7, antibodies blocking IFNAR-1 or IFNγR, or their corresponding isotype controls (500 µg/mouse) an hour after the LPS dose, and then challenged with ISO (300 mg/kg) on day 0. All mice were evaluated on day 1. **(A)** *Ifit1* and *Irf1* gene expression as evaluated by qPCR in isotype and antibody-treated mice (n=5-6/group). **(B)** Serum cardiac troponin I levels and heart weight-to-tibia length ratios. Flow cytometry analysis of **(C)** total $CD45^+$ cells, **(D)** $Ly6G^+$ neutrophils, $Ly6C^{hi}CD64^{lo}$ monocytes, $Ly6C^{lo}CD64^{hi}$ macrophages, $CD4^+$ T cells, $CD8a^+$ T cells, and $CD19^+$ B cells in the heart on day 1 (n=7-8/group). Data show mean ± SEM and were analyzed by unpaired, two-tailed t-test **(A-D)**, except for some that used the Mann-Whitney test **(**A, for anti-IFNγR versus isotype; D, for monocytes and macrophages, anti-IFNAR-1 versus isotype**)**. (Key *$P$<0.05, **$P$<0.01, ***$P$<0.001)

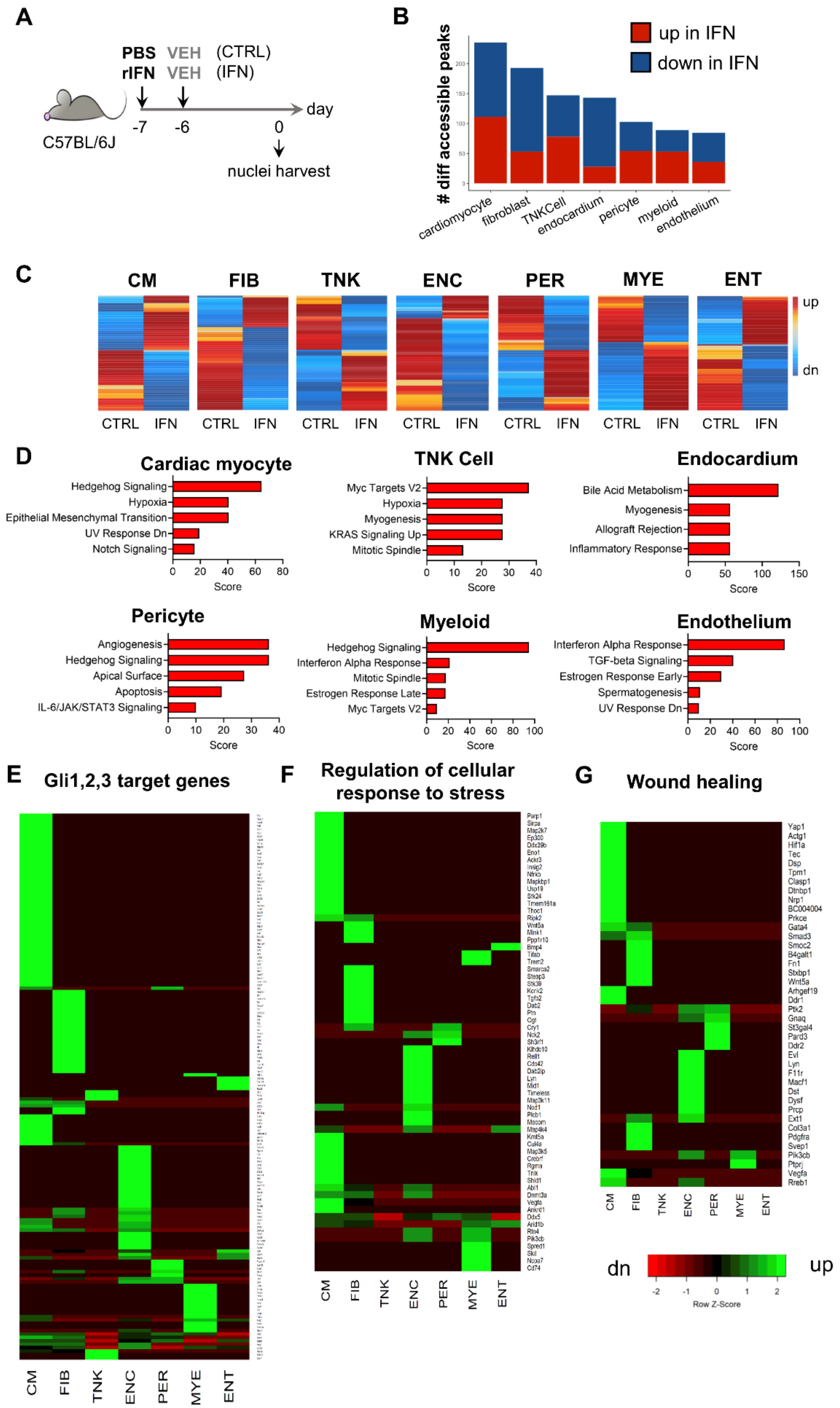


**Supplemental Figure S6. Multiomic analysis of recombinant interferon-treated hearts.**

**Supplemental Figure S6 (cont'd). (A)** Mice were injected i.p. with saline (PBS) on day -7 and vehicle (VEH) on day -6 (control, CTRL; same as used in **Figure 5**), or recombinant murine interferon-β1 (1 μg) and -γ (10 μg) (rIFN) on day -7 and vehicle (VEH) on day -6 (IFN), and nuclei were harvested from hearts on day 0 for multiomic snRNA-seq and snATAC-seq analysis (nuclei pooled from n=3/group). **(B)** Number of differentially accessible chromatin peaks in IFN compared to CTRL mice. **(C)** Heat maps of differentially accessible peaks across cardiac myocytes (CM), fibroblasts (FIB), TNK cells (TNK), endocardial cells (ENC), pericytes (PER), myeloid cells (MYE), and endothelial cells (ENT) in CTRL and IFN mice. **(D)** Pathways enriched from genes linked to chromatin peaks with increased accessibility in IFN hearts compared with CTRL hearts for predominant cell types in the heart. No enriched pathways were found for fibroblasts using the MSigDB Hallmark 2020 database. Heat maps of differentially expressed genes from the snRNA-seq across different cell types in IFN versus CTRL mice for those belonging to the **(E)** Gli1, 2, 3 transcription factor target genes (GTRD: P47806, Q0VGT2, Q61602), **(F)** regulation of cellular response to stress pathway (GO: 0080135), and **(G)** wound healing pathway (GO: 0042060).
